# Fine-tuning BERT models to extract transcriptional regulatory interactions of bacteria from biomedical literature

**DOI:** 10.1101/2024.02.19.581094

**Authors:** Alfredo Varela-Vega, Ali-Berenice Posada-Reyes, Carlos-Francisco Méndez-Cruz

## Abstract

Curation of biomedical literature has been the traditional approach to extract relevant biological knowledge; however, this is time-consuming and demanding. Recently, Large language models (LLMs) based on pre-trained transformers have addressed biomedical relation extraction tasks outperforming classical machine learning approaches. Nevertheless, LLMs have not been used for the extraction of transcriptional regulatory interactions between transcription factors and regulated elements (genes or operons) of bacteria, a first step to reconstruct a transcriptional regulatory network (TRN). These networks are incomplete or missing for many bacteria. We compared six state-of-the-art BERT architectures (BERT, BioBERT, BioLinkBERT, BioMegatron, BioRoBERTa, LUKE) for extracting this type of regulatory interactions. We fine-tuned 72 models to classify sentences in four categories: *activator*, *repressor*, *regulator*, and *no relation*. A dataset of 1562 sentences manually curated from literature of *Escherichia coli* was utilized. The best model of LUKE architecture obtained a relevant performance in the evaluation dataset (Precision: 0.8601, Recall: 0.8788, F1-Score Macro: 0.8685, MCC: 0.8163). An examination of model predictions revealed that the model learned different ways to express the regulatory effect. The model was applied to reconstruct a TRN of *Salmonella* Typhimurium using 264 complete articles. We were able to accurately reconstruct 82% of the network. A network analysis confirmed that the transcription factor PhoP regulated many genes (uppermost degree), some of them responsible for antimicrobial resistance. Our work is a starting point to address the limitations of curating regulatory interactions, especially for the reconstruction of TRNs of bacteria or diseases of biological interest.

## 1 Introduction

Every year, the number of scientific publications increases to such a magnitude that it is difficult to manually integrate, extract and organize the knowledge for subsequent analysis [1]. It was estimated that the annual growth rate of scientific literature in general is 4.10% and if this growth continues, in a period of 17.3 years the amount of literature will double [2]. For the biomedical field, it has been estimated that more than 3000 articles are published daily, a fact that represents an important barrier to access to the most relevant and updated knowledge for biologists and health professionals [3]. The manual extraction of biological information from scientific publications, named *literature curation*, has been the traditional way to extract, integrate and organize knowledge in biological databases [4]. As literature curation is demanding and time-consuming, machine learning and deep learning approaches of information extraction have been proposed for decades to support this task [5]. Technologies focused on the information extraction process are directed at a particular purpose, for example in the area of biomedical information, a goal is the extraction of different biological entities that help understand specific biological processes quickly and effectively in organisms or diseases [3]. Recently, Large language models (LLMs) based on pretrained transformers have gained interest as they have demonstrated surpassing the performance of classical machine learning approaches [6, 7]. Moreover, the benefit of fine-tuning a large pre-trained model with new limited data for a specific task (transfer learning) opens roads to address new problems of information extraction [8].

An example of a relevant problem is the reconstruction of biological networks from biomedical literature that has been a topic of interest since a while ago [9]. Here, we study the extraction of transcriptional regulatory interactions, which is the first stage to reconstruct a Transcriptional regulatory network (TRN) of bacteria. Despite the efforts to publish TRNs [10, 11], which are valuable resources organizing and integrating the knowledge of transcription regulation in bacteria, there is a lack of these networks, the existing ones are incomplete or they are not open access [12].

For our study, we defined a TRN as a set of transcriptional regulatory interactions between transcription factors and a regulated gen or a regulated operon (group of genes). These interactions play a leading role in the rapid response of bacteria to environmental signals and molecular stressors of the niche they colonize [13, 14]. The study of this kind of interactions may have an impact on relevant problems, such as antimicrobial resistance [15, 16]. An example in the bacterium *Escherichia coli* (*E. coli* ) is the study of the metabolism of glucose and diverse sources in relation to their expression of adjacent genes. Inquire this knowledge may help us to discover changes that may occur at the transcription level and may offer a new understanding of the metabolism in bacterial populations, influencing the processes of sugar absorption pathways, biotechnological processes and antibiotic optimization [17].

In a transcriptional regulatory interaction, if the regulatory effect is of the type *activation*, the transcription factor will promote the expression of the regulated gene. If the effect is of the type *repression*, the expression of the regulated gene will be inhibited by blocking the activity of the RNA polymerase [18, 19]. A regulatory interaction is therefore formed by a transcription factor, a regulated gene and an effect, which may be activation, repression, or simply *regulation*, when the effect exists, but the type of regulation is not reported.

Here, we fine-tuned six state-of-the-art architectures of Pre-training of Deep Bidirectional Transformers for Language Understanding (BERT) [8] to find the best model for extracting from biomedical literature the transcriptional regulatory interactions. To the best of our knowledge, the present study is the first effort to use BERT models to extract this specific kind of interactions. New standard benchmarks for biomedical relation extraction, such as BLUE, only includes drug-drug and chemical-protein interactions [20]. We employed techniques of *biomedical relation extraction*, a task of the Natural language processing (NLP) that aims at predicting from an input sentence whether two or more mentions of entities have some relationship (e.g., gene–disease, protein–protein, drug–drug) and, in some cases, the type of relation (i.e., cause, binding, induction) [5, 7, 21]. This task is basically a classification task, where a set of sentences are classified in a category [22]. A common strategy to address this task is to predict whether an interaction is true or false (binary classification) given a pair of mentions of entities and a sentence. For example, the mention of the Rob transcription factor and the mention of the *galT* gen in the following sentences must be classified as a true regulatory interaction:

- *”In this study, galT was the only gene repressed by Rob.”* [23]

In our work, we classified not only whether the regulatory interaction is true or false, we also classified the type of interaction in four categories (classes) depending on the type of effect of the transcription factor expressed over the regulated element: *activator*, *repressor*, *regulator*, and *no relation*. Therefore, our problem is defined as a multi-class classification task with exclusive categories. Examples of sentences of the three regulatory effects are shown in Table 1. Note that the way to express each type of effect is not necessarily done with verbs like *activate*, *repress* or *regulate*. Furthermore, the way to linguistically express the interaction is not always in active form ([TF] *activates* [GENE]). This makes our problem an interesting challenge for BERT architectures.

**Table 1.**
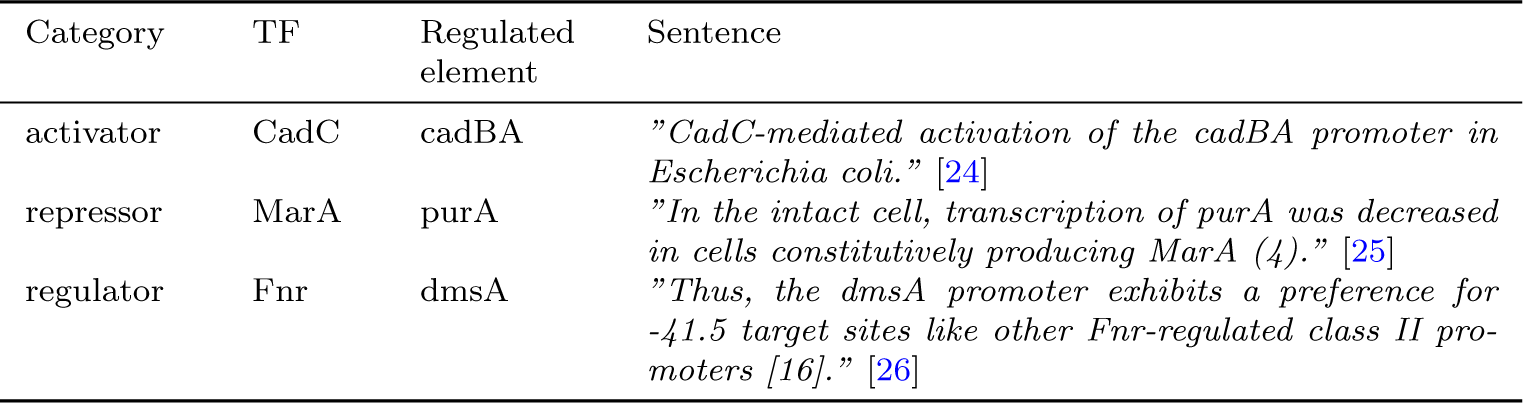
Examples of sentences expressing regulatory interactions between transcription factors (TF) and regulated elements.

We used a dataset of 1562 sentences of *E. coli* K-12 manually labeled in one of the four categories. This was randomly split in training (999 sentences), validation (250 sentences) and evaluation (313 sentences) datasets to fine-tune six state-of-the-art BERT architectures: BERT [8], BioBERT [22], BioLinkBERT [27], BioMegatron [28], BioRoBERTa [29] and LUKE [30]. Using the early stopping strategy when the crossentropy loss in the validation dataset did not improve in two epochs, we found a best model of LUKE architecture, which obtained cross entropy of 0.4024 and F1-Score Macro of 0.9107 measured in the validation dataset.

Our best LUKE model obtained a relevant final performance using the evaluation dataset: cross entropy of 0.4030, Precision of 0.8601, Recall of 0.8788, F1-Score Macro of 0.8685, and Matthew’s correlation coefficient of 0.8163. We found that the best classified category was *activator* (F1-Score Macro: 0.9082). The model addressed appropriately the imbalance of examples in categories of our dataset, as the second best classified category was the minority category *regulator*. The remaining categories achieved F-score higher than 0.82. From the analysis of the confusion matrix, we determined that the most confusing category was *no relation*. An examination of classification errors was carried out to explore the predictive capabilities of the best model. This examination revealed that the model learned different ways to express the regulatory effect, not only morphological variations, for example, *activated*, *activation* for the *activator* category, but also different lexical forms, for example, *enhance*, *stimulate*, *increase* for activation; and *inhibit* and *negative regulated* for the *repressor* category. The analysis of misclassified sentences revealed that some of them were correct predictions. The remaining errors were caused mainly by complex ways to express the interaction (auto-regulation, co-reference) or because the model highly weighted some words than others, an observation that we will investigate in future work using techniques of transformer interpretability.

Finally, our best model was applied for the reconstruction of a TRN of *Salmonella enterica* serovar Typhimurium using 264 complete articles. Utilizing a dataset of 3005 sentences with regulatory interactions extracted from the same articles by manual curation, we evaluated the reconstruction. The model was able to reconstruct 82% of the *Salmonella* TRN (Recall: 0.8217) demonstrating that our model may be used to extract regulatory interactions from literature of diverse bacteria. The network was visualized and analyzed with the Cytoscape system [31] and the PANTHER system [32]. We evaluated the degree value and reviewed the PhoP transcription factor (degree=180) in relation to its adjacent genes. We consider our work as a relevant starting point to address the limitations of access to biomedical knowledge, especially for the reconstruction of TRNs of different bacteria and diseases of biological interest.

## 2 Material and Methods

A general graphical scheme of the analysis carried out in the present work can be seen below (Figure 1) and will be explained in detail in the subsequent sections.

**Fig. 1.**
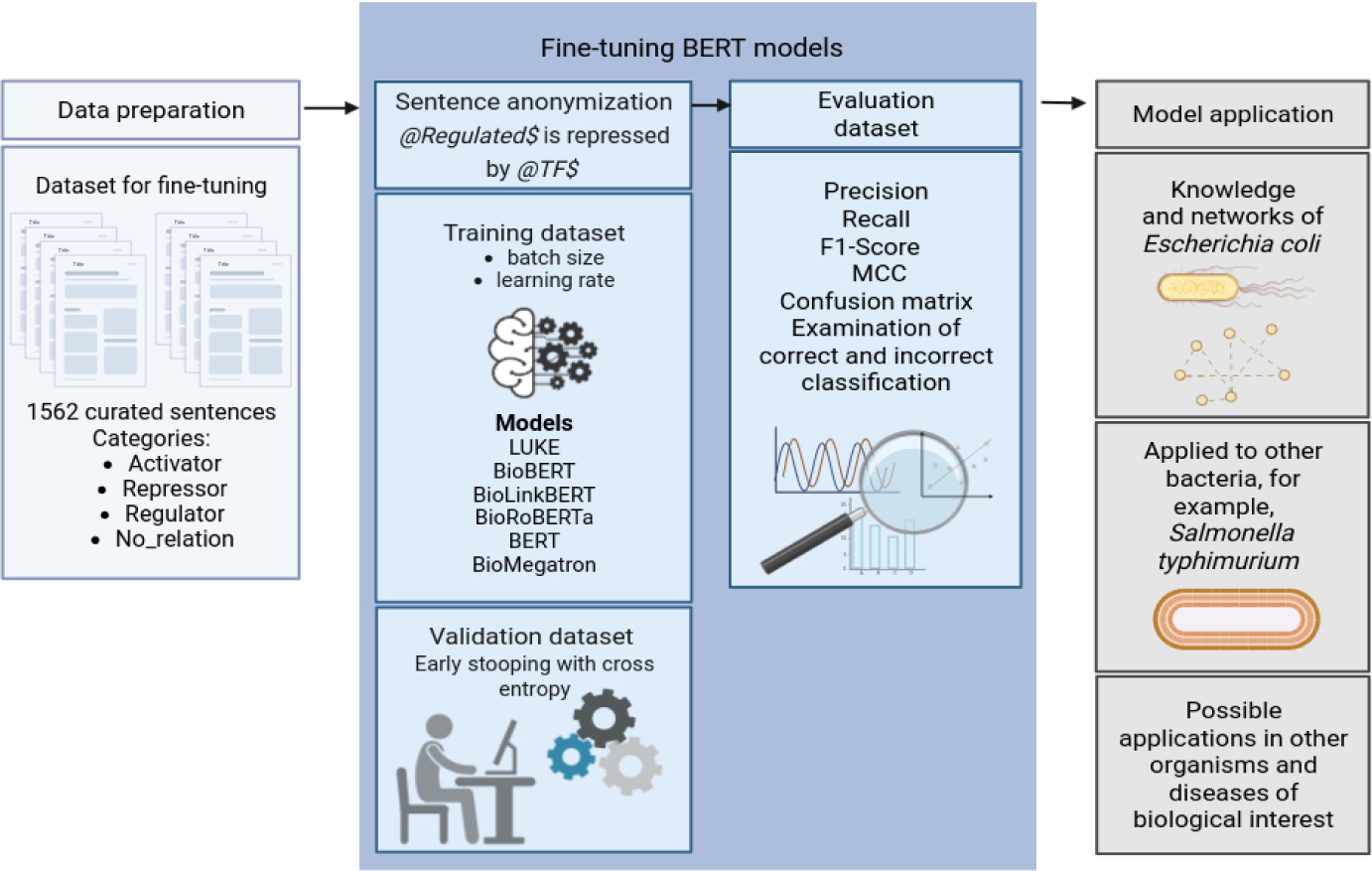
General graphical scheme of the analysis. Created with BioRender.com

### 2.1 Dataset for fine-tuning

RegulonDB is the database with the main transcriptional regulatory network publicly available of *E. coli* K-12 [10]. It comprises a large collection of regulatory interactions manually extracted for decades from biomedical literature. Some time ago, the curation team of RegulonDB compiled a set of 1083 sentences containing regulatory interaction using an assisted-curation strategy [33]. In this strategy, sentences containing the mention of at least one transcription factor, one gene/operon, and one regulatory keyword (e.g., *activates, regulation, inhibited, binding*) were filtered from articles using OntoGene/ODIN text mining tool [34]. Each sentence was manually examined to recover only those sentences that expressed a true regulatory interaction. Finally, the curator recorded the transcription factor, regulated element, sentence, and labeled the sentence with the regulatory effect: *activator*, *repressor*, or *regulator*. This dataset was provided by the curation team of RegulonDB.

A limitation of this initial dataset was that mentions of transcription factors and regulated elements (genes or operons) within the sentences were not textually marked (tagged). To fine-tune and evaluate a BERT model, it is required to know what mentions of the pair of entities (also named *target entities*) truly interact [22]. For instance, the initial dataset included the following sentence curated with the category *regulator*. However, we were not given the pair of mentions that actually have a regulatory interaction. Notice that the combination of the mention of the transcription factor RhaR and the first mention of the operon *rhaSR* clearly expresses the regulatory interaction (entities in in boldface); however, the combination of the mention of RhaR and the second mention of *rhaSR*, from a syntactic perspective, do not express any regulatory interaction (entities underlined):

- *”****<u>RhaR</u>*** *regulates transcription of* ***rhaSR*** *by binding promoter DNA spanning 32 to 82 relative to the <u>rhaSR</u> transcription start site”* [35].

Therefore, we curated the initial dataset of 1083 sentences to obtain a dataset suitable to fine-tune the BERT architectures. First, we recovered a list of transcription factors and regulated elements from the initial dataset. Using these lists, we separated those sentences with only one mention of each entity from those with multiple mentions either of the transcription factor, the regulated element or both. The latter were duplicated as many times as pairs of mentions and then curated to select the sentences with the pair of mentions expressing the true interaction (row 1 in Table 2). The remaining sentences were labeled with the category *no relation*, as they do not express a true interaction (row 2 in Table 2). The category *no relation* has been previously proposed for relation extraction datasets demonstrating that it is relevant for a model to improve its performance by learning from negative examples [36].

**Table 2.** Example of a sentence with two combinations of mentions (in boldface). First combination (row 1) expresses a true regulatory interaction between the RhaR transcription factor and the first mention of the *rhaSR* operon. Second combination (in row 2) does not express any regulatory interaction between RhaR and the second mention of *rhaSR*, then we labeled it with *no relation* category. The sentence was recovered from [35].

|  | Category | Sentence |
| --- | --- | --- |
| 1 | regulator | **RhaR** regulates transcription of **rhaSR** by binding promoter DNA spanning 32 to 82 relative to the *rhaSR* transcription start site |
| 2 | no_relation | **RhaR** regulates transcription of *rhaSR* by binding promoter DNA spanning 32 to 82 relative to the **rhaSR** transcription start site |

After such curation, we obtained a dataset with 1562 sentences to fine-tune the BERT architectures. The dataset included 66 transcription factors and 200 regulated elements. This dataset had an imbalanced distribution of sentences by category (Figure 2). The majority category was *activator* and the minority was *regulator*. The distribution of the length of the sentences (number of characters) was obtained in order to evaluate the heterogeneity and complexity of the dataset. The minimum sentence length was 27 characters and the maximum length was 665 characters (mean of 236, median of 223), the 75% of the sentences had a length less than 291 characters (Supplementary Figure S1). Sentences come from 119 different scientific articles (Supplementary Figure S2). The article that contributed the most sentences to the dataset gave the 6%. These characteristics reflect the richness and diversity of our dataset.

**Fig. 2.**
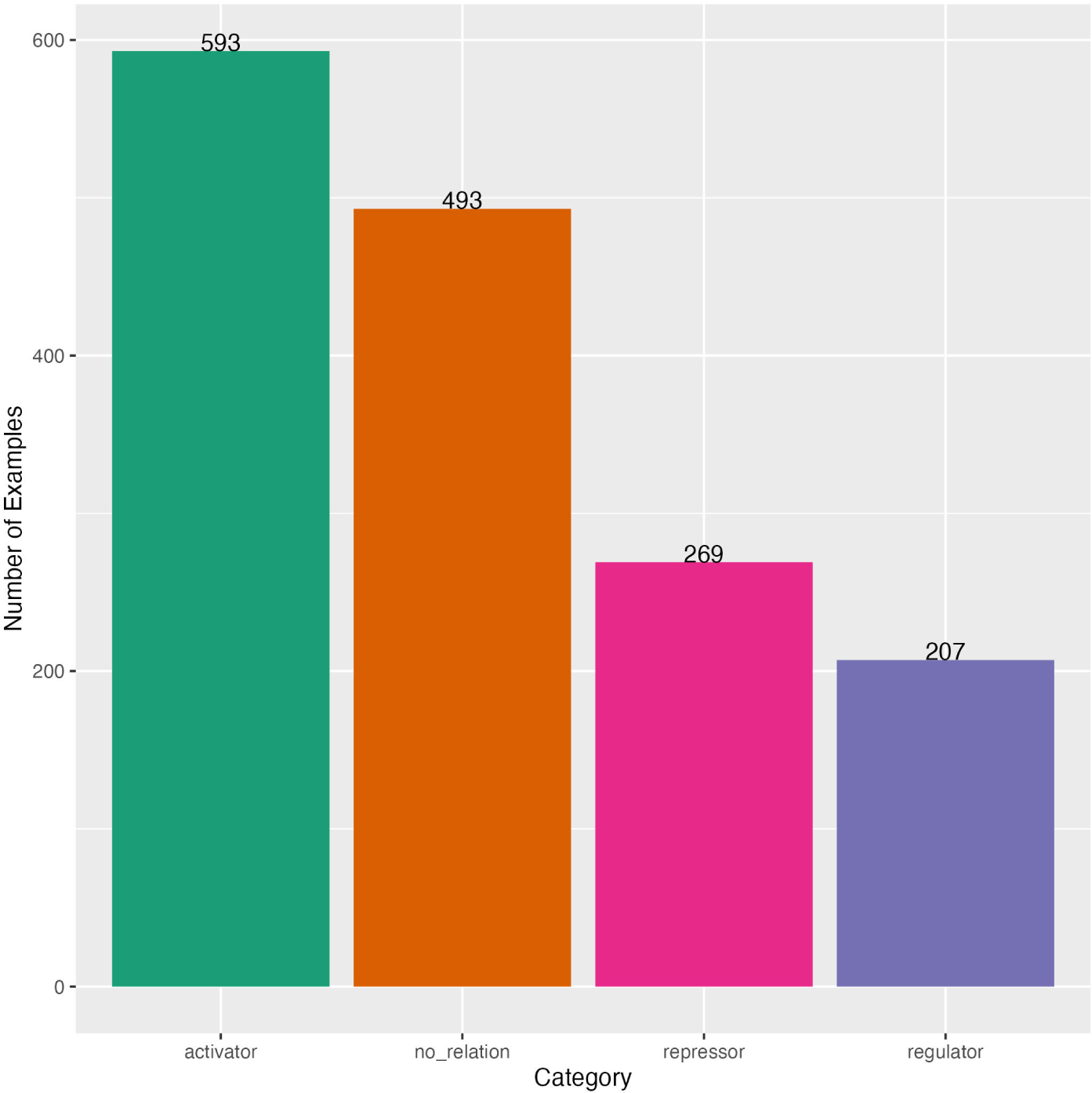
Category distribution in the dataset to fine-tune BERT architectures.

Our dataset was randomly split in 80% for fine-tuning and 20% for evaluation, then we split the fine-tuning dataset in 80% for training and 20% for validation. The final three datasets kept the same sentence distribution on categories as the complete dataset (Table 3).

**Table 3.** Distribution of sentences in the complete dataset and datasets for training, validation, and evaluation. Number of sentences followed by their percentage.

| Category | Complete | Training | Validation | Evaluation |
| --- | --- | --- | --- | --- |
| activator | 593 (38%) | 381 (38%) | 99 (40%) | 113 (36%) |
| no_relation | 493 (32%) | 316 (32%) | 77 (31%) | 100 (32%) |
| repressor | 269 (17%) | 169 (17%) | 44 (18%) | 56 (18%) |
| regulator | 207 (13%) | 133 (13%) | 30 (12%) | 44 (14%) |
|  | 1562 (100%) | 999 (100%) | 250 (100%) | 313 (100%) |

### 2.2 Dataset for model application

*Salmonella enterica* serovar Typhimurium (*Salmonella*) is one of the main pathogens that infect both humans and animals worldwide [37]. Transcriptional regulation of this bacterium has been studied for a while to face relevant problems, such as antimicrobial resistance [15]. The publication of a TRN for *Salmonella* has also received attention [11]. The curation team of RegulonDB also compiled a set of 3005 sentences containing regulatory interaction from 264 biomedical articles of *Salmonella* employing the same assisted-curation strategy described above. We utilized this dataset to evaluate the performance of our best model for reconstructing a TRN using complete articles. This dataset had the same limitation as the dataset of *E. coli* used for fine-tuning, that is, sentences did not have tags for the pair of entity mentions that truly participate in the interaction. Consequently, this dataset only had the three categories of regulation (*activator*, *repressor*, *regulator* ) and lacked the *no relation* category. Thus, the dataset contained the transcription factor, regulated element, sentence, and category.

The dataset included 91 unique transcription factors and 348 unique regulated elements. This had also an imbalanced distribution of sentences for each category (Figure 3). The majority category was *regulator* and the minority category was *repressor*. The minimum length of sentence (number of characters) was 23 characters and the maximum length was 2070 characters (mean of 201,median of 176), 75% of the sentences had a length less than 236 characters (Supplementary Figure S3). Sentences come from 264 different scientific articles (Supplementary Figure S4). The article that contributed the most sentences to the dataset was 8%. These characteristics are similar to those of the dataset used to find the best model and, at the same time, the distribution of the examples for each category is distinct enough to evaluate the performance of our best model during the model application phase (Supplementary Table S1).

**Fig. 3.**
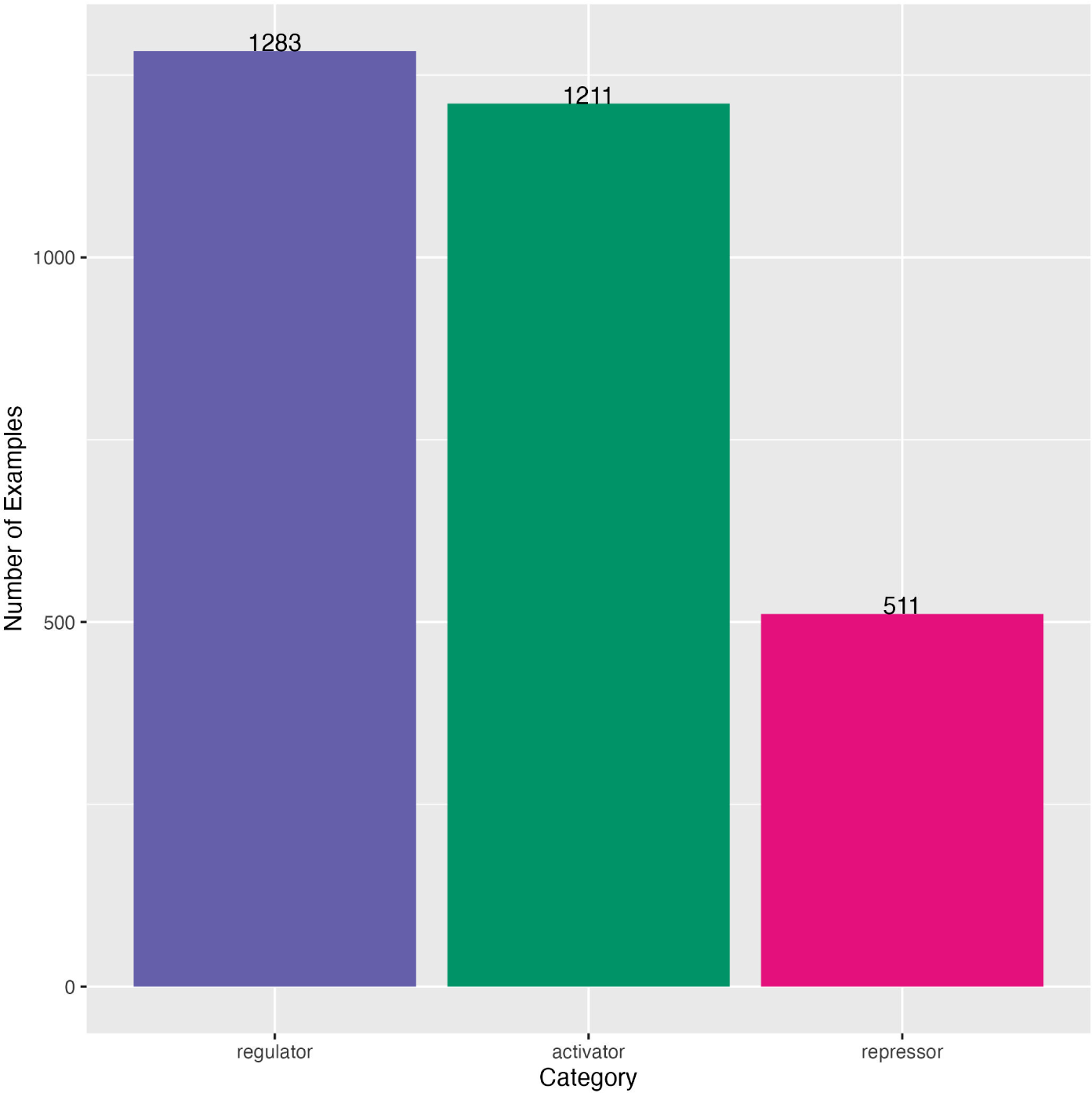
Category distribution in the dataset for model application.

### 2.3 BERT architectures

The *Bidirectional Encoder Representations from Transformers* (BERT) is a stateof-the-art large language model (LLM) based on the transformer architecture. Its capability to generate complex language representations lies in the two principal mechanisms that it inherits from the Transformer Encoder, as well as the massive pretraining over two subsets of text (*corpus*) [8]. These two main mechanisms are the multi-head self-attention and the position-wise fully connected feed-forward network, which, when assembled together, are able to create multiple representations of a word and its context within a sentence. The multi-head self-attention measures similarity scores between each word in the vocabulary by using positional encoding and word vector representations to predict the likelihood of the next occurring word or sentence [38]. Meanwhile, the second mechanism captures textual patterns in the training examples, with each value inducing a distribution over the output vocabulary that adds up to the next word or sentence prediction [39]. BERT was released to the public domain with the weights and biases learned during the pre-training phase using 800 million words from BookCorpus and 2500 million words from English Wikipedia. That was the turning point for more architectures to emerge using a transfer learning approach [40], taking BERT architecture and its previously acquired knowledge to fine-tune it for executing a specific language task, to excel in the understanding of a specialized domain, or simply to improve its overall performance.

In addition to BERT, we fine-tuned five more BERT models for our biomedical relation extraction task: BioBERT, BioLinkBERT, BioMegatron, BioRoBERTa and LUKE. The *Bidirectional Encoder Representations from Transformers for Biomedical Text Mining* (BioBERT) is a domain-specific model pre-trained on 4.5 billion words from PubMed abstracts and 13.5 billion words from PubMed Central full-text articles [22]. BioLinkBERT is a model that incorporates link knowledge by pre-training the architecture with hyperlinks between Wikipedia articles and citation links (references) from 21 GB of PubMed articles [27]. BioMegatron is an architecture designed to extend the number of trainable parameters of BERT beyond 345 million by implementing efficient model parallelism. Moreover, BioMegatron extends the pre-training corpus by adding 4.5 billion words from PubMed abstracts and 1.6 billion words of a CC0licensed Commercial Use Collection of the PMC full-text corpus [28]. BioRoBERTa is the biomedical domain version of the *Robustly Optimized BERT Approach* (RoBERTa) model, which pointed out that BERT was under-trained and should be trained longer, using bigger batches, longer sequences and dynamically changing the masking pattern during training. Furthermore, it was added 76 GB of English news articles, 38 GB of open web text content, and 31 GB of a dataset matching story-like style of Winograd schemas [41]. For the biological domain version, BioRoBERTa uses 2.68 million fulltext papers from The Semantic Scholar Open Research Corpus [29]. The *Language Understanding with Knowledge-based Embeddings* (LUKE) makes adjustments to the training following the RoBERTa architecture. It also implements a new pre-training task, predicting randomly masked words and entities in a large entity-annotated corpus retrieved from Wikipedia. Moreover, a slight change in the self-attention mechanism was done to compute attention scores based on the type of token, creating the entity-aware self-attention mechanism [30].

### 2.4 Experimental design

#### 2.4.1 Fine-tuning

To fine-tune models of BERT, BioBERT, BioLinkBERT, BioMegatron and BioRoBERTa architectures for relation extraction tasks, entity mentions must be anonymized [22]. Therefore, in each of the 1562 sentences, the mentions of the transcription factor and the regulated element were anonymized using the @TF$ and @Regulated$ pre-defined tags, respectively (Table 4). This is a common procedure to prepare sentences for fine-tuning BERT models [20, 21, 42]. The sentences with anonymized entities were input to the tokenizer of each architecture, which generated a numerical representation of each word so that the model could be fine-tuned. In the particular case of the LUKE architecture, the start and the end positions of each entity within the sentence (spans) are required by its tokenizer, which subsequently generates the entity masking. Then, the sentence and the entity spans were input to LUKE architecture (Table 4).

**Table 4.** Examples of inputs to BERT architectures for the sentence: *marA expression is repressed by MarR*. The BERT, BioBERT, BioLinkBERT, BioMegatron and BioRoBERTa required the sentence with anonymized entity mentions. LUKE required the original sentence and the entity spans.

| Architecture | Input |
| --- | --- |
| BERT, BioBERT, BioLinkBERT, BioMegatron, BioRoBERTa | @Regulated\$ expression is repressed by @TF\$ |
| LUKE | *marA expression is repressed by MarR* [[0, 4], [78, 82]] |

To find the best BERT model, we used the strategy of grid search for the batch size and the learning rate based on the early stopping when the cross entropy does not improve in two epochs using the validation dataset. This strategy is commonly utilized to train Deep learning models [43]. Briefly, training a Deep learning model consists of searching for millions of weights (parameters) which generates the smallest difference between the predictions provided by the model and the true categories of training data. One way to measure this difference is through a loss function. For multi-class classification problems, a common way to calculate this function is by means of the cross-entropy loss function [44, 45]. This function takes the output from the softmax activation function (the probability that an example belongs to each category), and calculates a scalar value quantifying the difference (error) between the predicted probability distribution *p* and the true categories *y*:

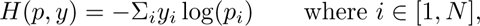

where *p* is a vector of probabilities from the softmax function of size *N* , that is, the number of categories; and *y* is usually a binary vector of size *N* , where 1 indicates the true category and 0 the false one. For this function, the worst value is +∞ and the best value is 0.

Finding the parameters that allow us to accurately predict the categories of the training data is important, but a characteristic that gives robustness to a model is the ability to perform correctly with new data (different from those used in training). This generalization capacity can be achieved with the so-called regularization techniques. Early stopping is one of these techniques that is also useful for saving computational resources because the model stops its training when the metric to be optimized does not improve in a certain number of cycles. The patience is the number of cycles before stopping the training and it should be a low value when the input data is few and very informative. In the present work, for the early stopping we calculated the cross entropy in the validation dataset (*patience* = 2) as it has been recommended in previous works [43].

The size of the adjustment of weights is called learning rate (lr) and the algorithm that uses this value to calculate and update the weights is called optimizer [45]. We used the AdamW optimizer for all experiments. The advantage of using this is that it implements an improvement in the regularization of its base algorithm Adam (adaptive moment estimation) [46]. AdamW makes the weight decay independent, that is, it changes the magnitude of the weights by favoring small values regardless of the direction in which the parameters will be updated [47].

We also used the mini-batches strategy to get the best BERT model. In this strategy, a dataset is divided into subsets named batches, so the number of subsets is equal to the number of examples divided by the batch size. The model goes through each batch (step) and, at the end of each batch, it updates the weights using the optimizer algorithm. When all the batches have been covered, it is said that an epoch has passed [48].

To conduct a fair compassion of models, we use the same grid of values for the batch size and the learning rate in all experiments. We selected these values from those commonly recommended for fine-tuning in the original articles of the six architectures:

- batch size: 10, 16, 32, 64
- learning rate: 1e-5, 3e-5, 3e-5

The experimental grid resulted in 72 fine-tuned models, twelve for each architecture. We selected the best hyper-parameters (best model) using the lowest cross entropy in the validation dataset. Pre-trained BERT models were downloaded from *Hugging Face* (https://huggingface.co/), details are described in Supplementary Table S2. We coded Python scripts (version 3.11.3) for the fine-tuning, evaluation and inference using the specialized framework for deep learning *PyTorch Lightning* (https://lightning.ai/docs/pytorch/stable/). A list of specialized libraries employed in our study are detailed in Supplementary Table S3. All BERT models were fine-tuned in CPUs (Linux CentOS 4.18.0-499.el8.x86 64, 40 CPUs).

#### 2.4.2 Evaluation

The final best model was obtained by retraining the best architecture using the best hyper-parameters and the total sentences from the training and the validation datasets together. We evaluated this final model using the evaluation dataset, so we measured the model generalization, that is, the performance of the model to classify sentences in new datasets. We calculated several metrics: precision, recall (sensitivity), F1-Score Macro, and Matthew’s correlation coefficient [49]. To expand the understanding of the predictive capabilities of the model, we examined the confusion matrix, the metrics for each category, and the correct and incorrect predictions on the evaluation dataset.

For a binary classification problem with the *positive* and *negative* categories, the precision score is the fraction of examples predicted correctly by the model:

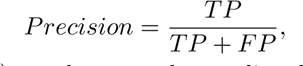

where *TP* (true positives) are the examples predicted as positive category that are actually true; and *FP* (false positives) are the examples predicted as positive category that actually correspond to the negative category. The recall is the fraction of correctly predicted examples of a category among the total examples of that category:

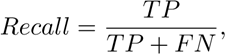

where *FN* (false negatives) are the examples incorrectly predicted as the negative category. The F1-Score is the harmonic mean of Precision and Recall:

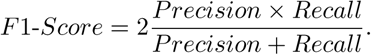

We used F1-Score Macro, which calculates the statistics for each label and averages them, attributing equal importance to all classes despite the imbalance of the categories. For these three metrics 0 is the worst value and 1 is the best value. The Matthew’s correlation coefficient (MCC) measures the correlation of the true categories *c* with the predicted categories *l*; the worst value is −1, and the best value is 1. This metric is recommended for classification problem with imbalance categories, as it gives a high score only if the classifier correctly predicted most of the categories [50]:

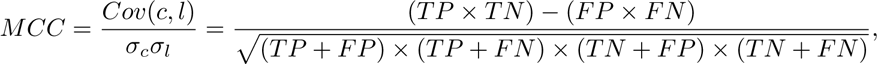

where *Cov*(*c, l*) is the covariance of the true categories *c* and the predicted categories *l*; *σ_c_* and *σ_l_* are the standard deviations, respectively. The *TN* (true negatives) are the correctly predicted examples of the negative category.

A confusion matrix is relevant to observe the performance of a classifier for a multiclass task, because it shows for each pair of classes ⟨*c*1*, c*2⟩ how many examples of category *c*1 are incorrectly classified as category *c*2, and vice versa. This matrix allows to point out opportunities to improve the performance of a classifier [49], as we may find the more confusing category:

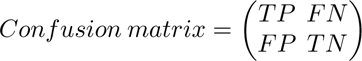

### 2.5 Model application

We applied our final best model to reconstruct a TRN of *Salmonella* using complete biomedical articles. The idea is to explore the performance of the model to deal with the reconstruction of TRNs of different bacteria, which is one of our longterm goals. To evaluate the *Salmonella* TRN reconstruction, we used the dataset with 3005 curated sentences from 264 articles (section 2.2); however, instead of measuring the classification of theses sentences, we measured the capability of the model to extract the regulatory interactions (transcription factor, regulated object and category) reported in the curated dataset. In other words, we compared the regulatory interactions extracted by the model with the regulatory interactions extracted by the curator. A complete study using large TRNs publicly available will be done in future work. This change in evaluation strategy is mainly due to the fact that to evaluate our method with a published TRN (for example, RegulonDB) we will not have the sentences from which the interactions were curated, but only the set of interactions.

Then, we collected the 264 PDF files. For articles without open-access rights, downloading rights were granted by our institution. PDF files were converted into plain text using an in-house developed tool. Sentence split and tokenization of sentences were performed with the Stanford CoreNLP tool [51]. To prepare the sentences for LUKE (the best model), we obtained a list of transcription factors and a list of regulated elements from the curated dataset of *Salmonella*. Then, we automatically obtained the spans of the pair of entity mentions by searching the entities from the lists within each sentence of the 264 articles. From the total sentences of the articles, 14349 had at least one transcription factor and one regulated element. These sentences and the spans were input to the best model to predict the category (model inference).

The classified sentences were filtered to discard those predicted with *no relation* category. From the filtered sentences, we obtained the unique regulatory interactions, which were searched in the curated dataset to assess the performance of the model. Our evaluation was in two ways. First, we performed an evaluation by comparing the complete interaction, that is, using the three elements: transcription factor, regulated element and category. Second, we evaluated only the interacting entities, that is, only the regulator and the regulated element. For both types of evaluations, repetition of sentences expressing the same regulatory interaction could benefit the model.

We calculate Precision, Recall, and F1-Score. The MCC was not calculated, as we did not have negative examples (false regulatory interactions) in the curated dataset and the number of true negatives could not be obtained. The metrics were calculated the same as for the evaluation of the best model (see section 2.4.2); however, in this case, the true positives (*TP* ) were the extracted regulatory interactions that were present in the curated dataset; the false positives (*FP* ) were the extracted regulatory interactions that were not present in the curated dataset; and the false negatives (*FN* ) were the regulatory interactions from the curated dataset that were not extracted by the model [49]. To get a better understanding of the model performance to reconstruct the TRN, we manually examined a sample of incorrect predictions (false positive cases).

Finally, the regulatory interactions extracted by the model were visualized and analyzed with Cytoscape (https://cytoscape.org/). Using this software, we calculate some network centrality measurements (betweenness centrality, degree distribution). The transcription factor with the highest connectivity in the network (degree) was considered along with its community (connected regulated elements). The list of regulated elements was entered into the PANTHER system to identify their biological processes, functional molecular characteristics and an analysis of overrepresentation test Fisher’s exact (https://pantherdb.org/).

## 3 Results

### 3.1 Performance of the best model

To select the hyper-parameter combination that maximized the performance of each BERT architecture, we trained with the training dataset and evaluated using the validation dataset. As we mentioned, we used an early stopping strategy (*patience* = 2) based on cross entropy. The summary of the hyper-parameters found for the best model of each architecture, as well as the metrics, are shown in Table 5; best result in boldface. The best model was from LUKE architecture with a batch size of 32, a learning rate of 0.00001, trained during 12 epochs and 415 steps. With these hyper-parameters LUKE reached an F1-Score Macro of 0.9107 and a loss of 0.4024.

**Table 5.** The best hyperparameters for each model evaluated over the validation dataset. Best result in boldface.

| Model | F1-Score<br>Macro | Loss | Epoch | Steps | Batch<br>size | Learning<br>rate | Runtime |
| --- | --- | --- | --- | --- | --- | --- | --- |
| **LUKE** | **0.9107** | **0.4024** | **12** | **415** | **32** | **0.00001** | **6h 55m 34s** |
| BioBERT | 0.8983 | 0.4633 | 6 | 111 | 64 | 0.00005 | 9h 38m 14s |
| BioLinkBERT | 0.8893 | 0.4137 | 7 | 503 | 16 | 0.00001 | 3h 58m 20s |
| BioRoBERTa | 0.871 | 0.4547 | 10 | 351 | 32 | 0.00001 | 5h 3m 34s |
| BERT | 0.8552 | 0.5319 | 7 | 503 | 16 | 0.00003 | 4h 3m 5s |
| BioMegatron | 0.8434 | 0.8096 | 6 | 440 | 16 | 0.00005 | 10h 29s |

A graphic summary of the results obtained by the twelve LUKE training runs can be seen in Figure 4, figures of remaining architectures are shown in Supplementary material (Figures S5-S9). These figures were obtained using *Weights & Biases* (https://wandb.ai/site), a platform designed for the organization and development of artificial intelligence workflows. Each line in the figure represents a combination of hyper-parameter values (*Batch size*, *Learning rate*, *Epoch*) and for each combination the cross entropy loss value (*Validation Loss*) and F1-Score Macro (*Validation f1 epoch*) obtained on the validation dataset is shown. We did not observe any tendency between the combinations of hyper-parameter values and loss values. An interesting case was BioBERT, as the majority of combinations of the hyper-parameter values obtained close loss values (Supplementary Figure S6).

**Fig. 4.**
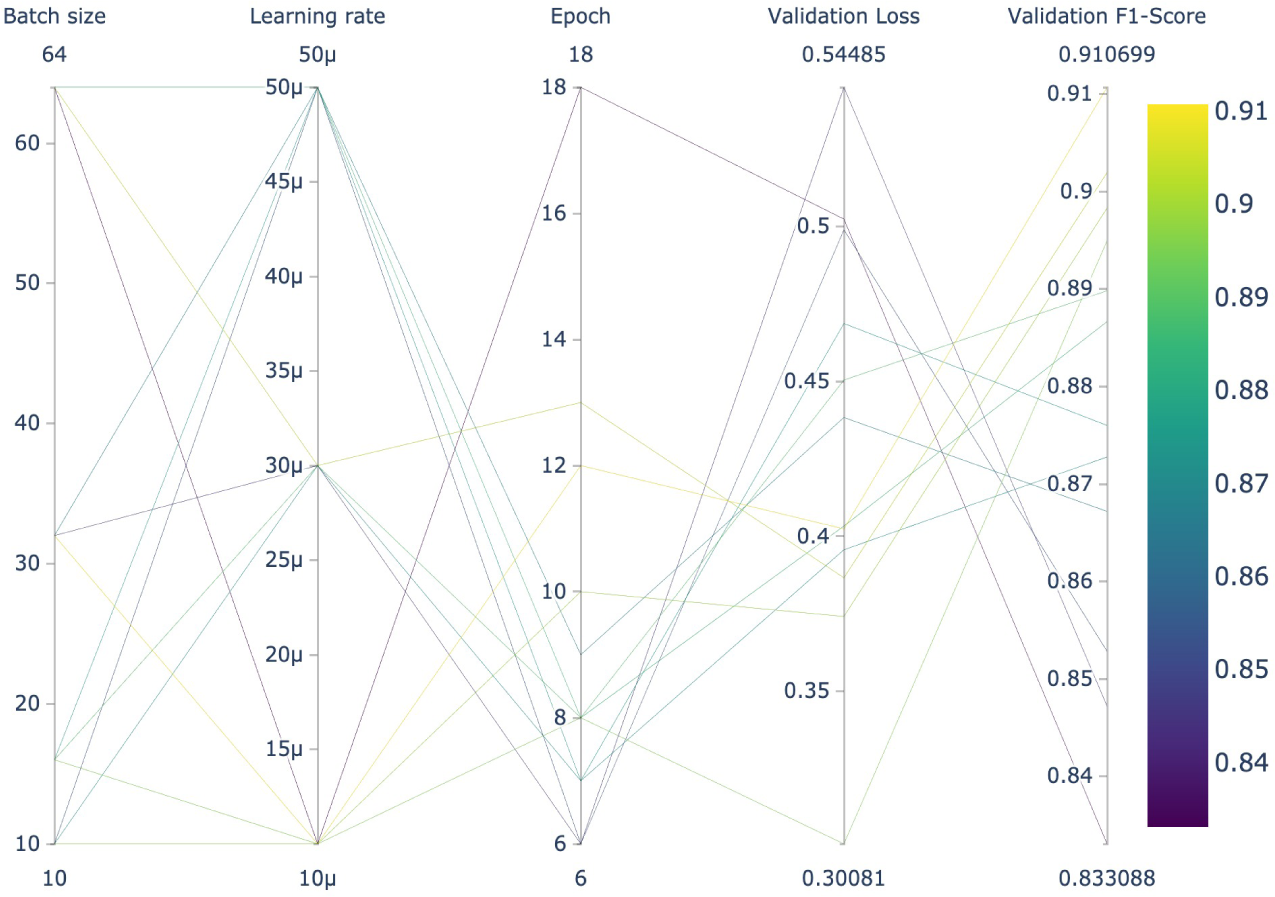
Hyper-parameter search of the twelve LUKE models evaluated on the validation dataset.

On the evaluation step using the evaluation dataset, the final best LUKE model also obtained relevant metrics (Table 6). This model achieved balanced scores of Precision (0.8601) and Recall (0.8788), which led to a final F1-Score Macro of 0.8685. The MCC was also high (0.8163), showing that the model dealt well with the imbalance of sentences in the categories. The decrease in value of the F1-Score obtained in the fine-tuning step against that obtained in the evaluation step shows that the model was overfitted; a common result for complex deep learning architectures such as transformers. This limitation will be addressed in future work by adopting some additional strategies proposed in literature [52, 53].

**Table 6.**
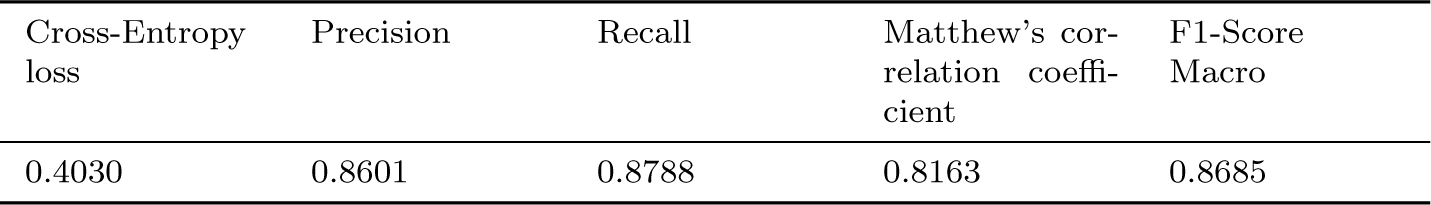
Performance metrics of the final best LUKE model obtained on the evaluation step using the evaluation dataset.

| Cross-Entropy loss | Precision | Recall | Matthew’s correlation coefficient | F1-Score Macro |
| --- | --- | --- | --- | --- |
| 0.4030 | 0.8601 | 0.8788 | 0.8163 | 0.8685 |

The best classified categories were *activator* (F1-Score Macro: 0.9282) and *regulator* (F1-Score Macro: 0.8915), and all categories obtained an F1-Score Macro up to 0.8200 (Table 7). Notice that *regulator* was the minority category (support = number of sentences), which confirmed that our model performed well despite the imbalance of sentences in categories.

**Table 7.**
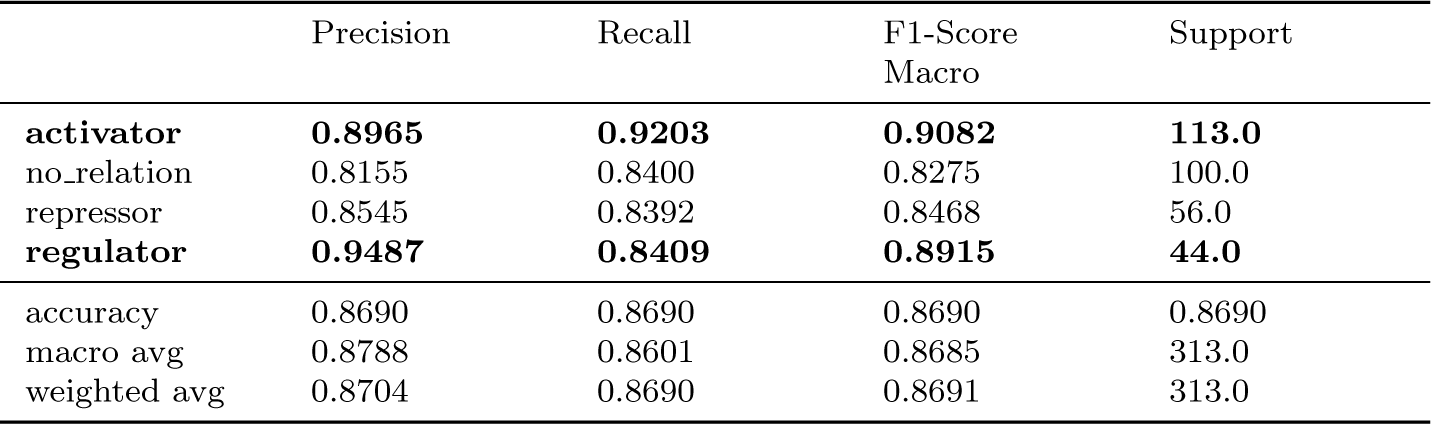
Classification report of the best LUKE model obtained on the evaluation step.

Finally, we report here the confusion matrix obtained by our best model. This matrix deeply depicts the classification results for each category with emphasis on how the model confuses them. The analysis of this matrix is presented in the Discussion (section 4).

### 3.2 *Salmonella* TRN reconstruction

We classified with our best LUKE model the 14349 sentences obtained from the 264 articles of *Salmonella*. These sentences had at least one mention of a transcription factor and one mention of a regulated element (see section 2.5). From the classified sentences, we discarded those with *no relation* category obtaining 8256 sentences. As we want to reconstruct a TRN, we get unique regulatory interactions, that is, a combination of transcription factor, regulated element and category. Then, 1826 unique interactions were recovered, including 90 unique transcription factors and 324 unique regulated elements.

To evaluate the TRN reconstruction, we obtained the unique regulatory interactions from the curated dataset of *Salmonella* (see section 2.2). We identified 909 unique regulatory interactions, which contained 91 unique transcription factors and 348 unique regulated elements. We observed a very similar distribution of interactions between curated and predicted categories (Supplementary Table S4). We performed a second evaluation with the combination of only the transcription factor and the regulated element (interacting entities). In this case, 1031 unique predicted interactions and 641 unique curated interactions were obtained from the curated collection.

We summarize the results of the two evaluations in Table 8. Our best model obtained a relevant recall recovering the curated complete interactions (0.8217). Nevertheless, the precision score of the model was low (0.4090) due to the high number of false positives (extracted interactions that did not appear in curated interactions). Note that in fact many of these might well be correctly predicted interactions absent in the curated collection. The evaluation with only interacting entities exhibited the same performance of Precision and Recall scores. We observed that discarding the type of effect, our best model was able to recover the 87% of the regulatory interactions (F1-Score: 0.8720); however, these predictions do not predict the regulatory effect.

**Table 8.** Evaluation of *Salmonella* TRN reconstruction with our best model. Evaluation with complete interaction refers to evaluating the correct extraction of the combination of transcription factor, regulated element and category, whereas evaluation with interacting entities refers to evaluating the correct extraction of the combination of transcription factor and regulated element.

|  | Evaluation with complete interaction | Evaluation with interacting entities |
| --- | --- | --- |
| True (curated) interactions | 909 | 641 |
| Extracted interactions | 1826 | 1031 |
| True positives | 747 | 559 |
| False negative | 162 | 82 |
| False positive | 1079 | 472 |
| Precision | 0.4090 | 0.5421 |
| Recall | 0.8217 | 0.8720 |
| F1-Score | 0.5462 | 0.6686 |

We achieved a network analysis of the reconstructed TRN of *Salmonella*. The TRN showed 414 nodes (transcription factors and regulated elements) and 1826 edges (interactions). A visualization of the TRN is available in Supplementary Figure S10. The PhoP transcription factor has the highest value of degree (180) and the fifth highest value of betweenness centrality (0.2642). Betweenness centrality measures the relevance of a node in terms of shortest paths in the network [54]. Nodes with the highest betweenness centrality may be taken as relevant links (bridges) between nodes in a network. The degree is the number of edges connected to the node [54]. In a TRN, nodes with the highest degree are transcription factors regulating many regulated entities. The complete output of the Cytoscape network analysis for degree and betweenness centrality is available in Supplementary Table S7.

As PhoP attained the highest degree, we selected its community (connected regulated entities) for visualization and analysis. In this community network, 99 nodes with 420 edges were identified. The community network was visualized with the Cytoscape system (Figure 6). The PhoP’s adjacent genes were analyzed for their molecular function and biological processes using the PANTHER system. Molecular function was described as follows: ATP-dependent activity (GO:0140657) with 3.3% and 2.8%, antioxidant activity (GO:0016209) with 1.6% and 1.4%, binding (GO:0005488) 11.5% and 9.7%, catalytic activity (GO:0003824) 13.1% and 11.1%, molecular transducer activity (GO:0060089) 9.8% and 8.3%, transporter activity (GO:0005215) 4.9% and 4.2%. The biological processes data were: response to stimulus (GO:0050896) with 3.3% and 2.9%, cellular process (GO:0009987) 8.2% and 7.4%, metabolic process (GO:0008152) 9.8%, and 8.8%, biological regulation (GO:0065007) 9.8% and 8.8%.

Finally, an over-representation analysis was carried out. The Fisher’s exact test described two biological processes: Magnesium ion transmembrane transport has raw *P* value of 7.51e*^−^*^05^ and FDR of 2.24e*^−^*^02^, and Phosphorelay signal transduction system with raw *P* value of 1.19 e*^−^*^06^ and FDR of 9.42e*^−^*^04^. By comparison, we performed the same over-representation analysis using the curated interactions for PhoP, which comprises 72 regulated elements. We found that the same biological processes were enriched: Magnesium ion transmembrane transport (raw *P* value of 3.03e*^−^*^05^ and FDR of 1.44 e*^−^*^02^) and Phosphorelay signal transduction system (1.20e*^−^*^05^ and FDR of 1.43e*^−^*^02^).

## 4 Discussion

In the information extraction field using BERT models, biomedical relation extraction is proposed as a classification problem: given a sentence where the mentions of entities are anonymized, the model has to predict if the entities truly interact or not (binary classification). Another approach is asking the model to predict the type of interaction (multi-class classification). Here, we compared six BERT architectures and selected the best model to classify the type of regulatory effect given a sentence and a pair of mentions of a transcription factor and a regulated element (gene or operon), based on curated knowledge obtained from RegulonDB and curators. Afterwards, we applied our best model to reconstruct a TRN of *Salmonella* using 264 complete articles and we evaluated this reconstruction using a set of sentences curated from the same article collection. The discussion of results are presented in the following sections.

### 4.1 Our model accurately extracts regulatory interactions

Our best model, based on LUKE architecture, obtained a balanced performance of classification observed in the F1-Score Macro of 0.8685 (Table 7). The Precision score (0.8601) shows that, on average, when the model predicts a category, 86% of the times the prediction is correct. The Recall score (0.8788) indicates that, on average, the model correctly classifies 87% of all input sentences. According to the Matthew’s correlation coefficient (0.8163), our model has a relevant performance despite the imbalance of sentences in the categories; this score indicates that our model shows a strong positive correlation between its predictions and the true categories of the sentences, that is, the predictions are not random. We consider that the high performance of LUKE is due to the fact that its architecture is based on the optimization of RoBERTa and in the pre-training step the entities and words are independently processed by an entity-aware self-attention mechanism. This capability to represent entities distinctly from the rest of the words was relevant in our classification task to detect regulatory interactions.

It is noteworthy that LUKE outperformed specialized transformers pretrained with biomedical literature (BioBERT, BioLinkBERT, BioMegatron and BioRoBERTa). This may indicate that it is as important to pre-train a BERT model using a domain-specific corpus as it is to pre-train the model for a specific task, so the model would learn the structure and patterns found in both specialized and nonspecialized languages. As far as we know, LUKE has not been previously evaluated for biomedical relation extraction.

From the analysis of the performance for individual categories (Table 7), we observed that the best classified category was *activator* (F1-Score Macro: 0.9082). In addition, it was the top recovered category (Recall: 0.9203) and the second best in precision (0.8915). It is interesting that the second best predicted category was *regulator*, although it was also the minority category in the evaluation dataset (44 examples); this fact confirms the capacity of our model to deal with category imbalance. Furthermore, the *regulator* category was the class with the highest precision (0.9487), showing that our model is reliable when predicting this category. In general, all the categories obtained a relevant F1-Score (above 0.82) regardless of the number of examples (support) of each category.

The examination of the confusion matrix obtained from the predictions in the evaluation dataset allows us to observe which categories our model confuses the most and which the least (Figure 5). The best model learned to efficiently differentiate among the three categories of regulation (*activator*, *repressor* and *regulator* ). Only one sentence that was actually *regulator* and one sentence that was actually *repressor* was misclassified as *activator* (first column of the confusion matrix). The third column in the confusion matrix shows the sentences predicted as the *regulator* category, the model misclassified two sentences, one of *activator* and one of *repressor*. The model made errors predicting the category *repressor* by misclassifying two sentences of *activator* category, but it did not confuse the *repressor* category with the *regulator* category (fourth column in the confusion matrix).

**Fig. 5.**
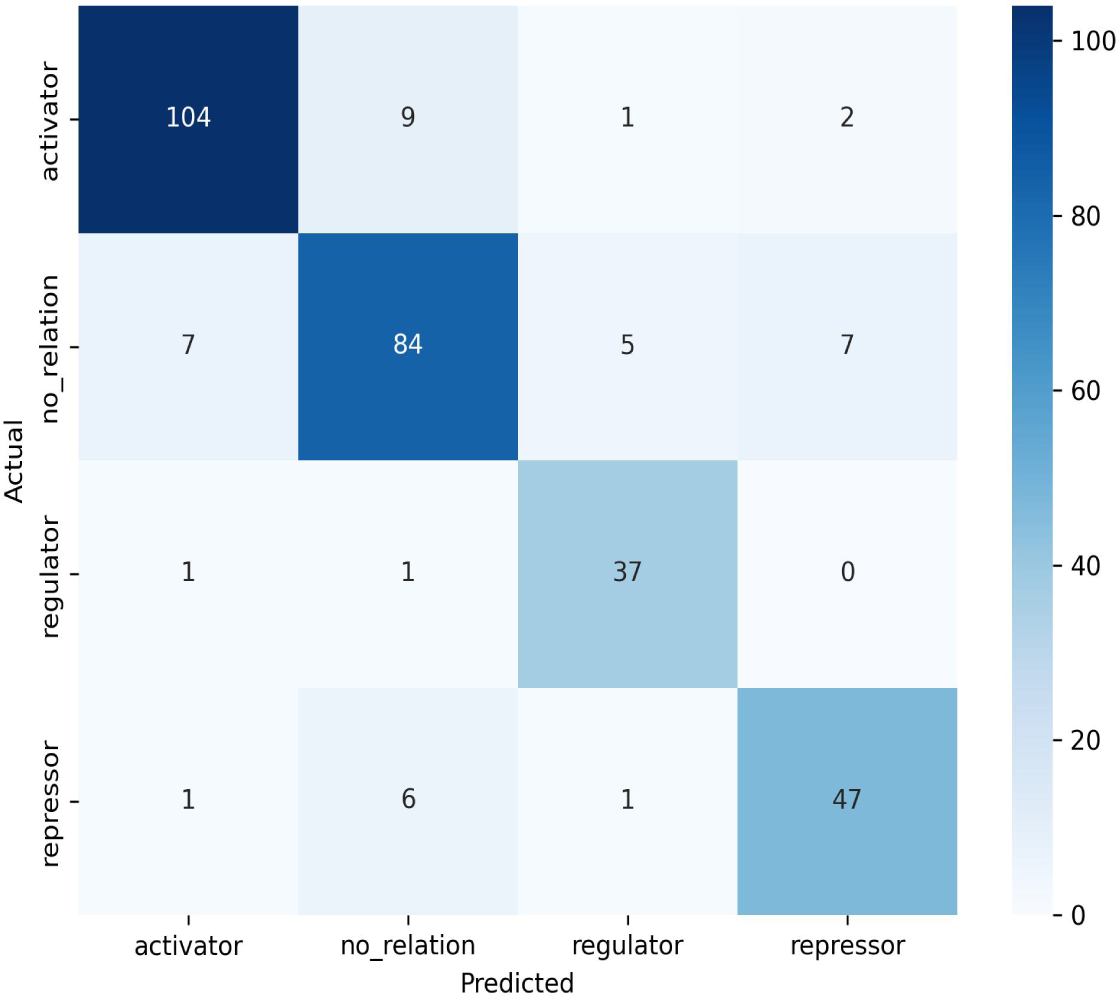
Confusion matrix of the best LUKE model obtained on the evaluation step.

**Fig. 6.**
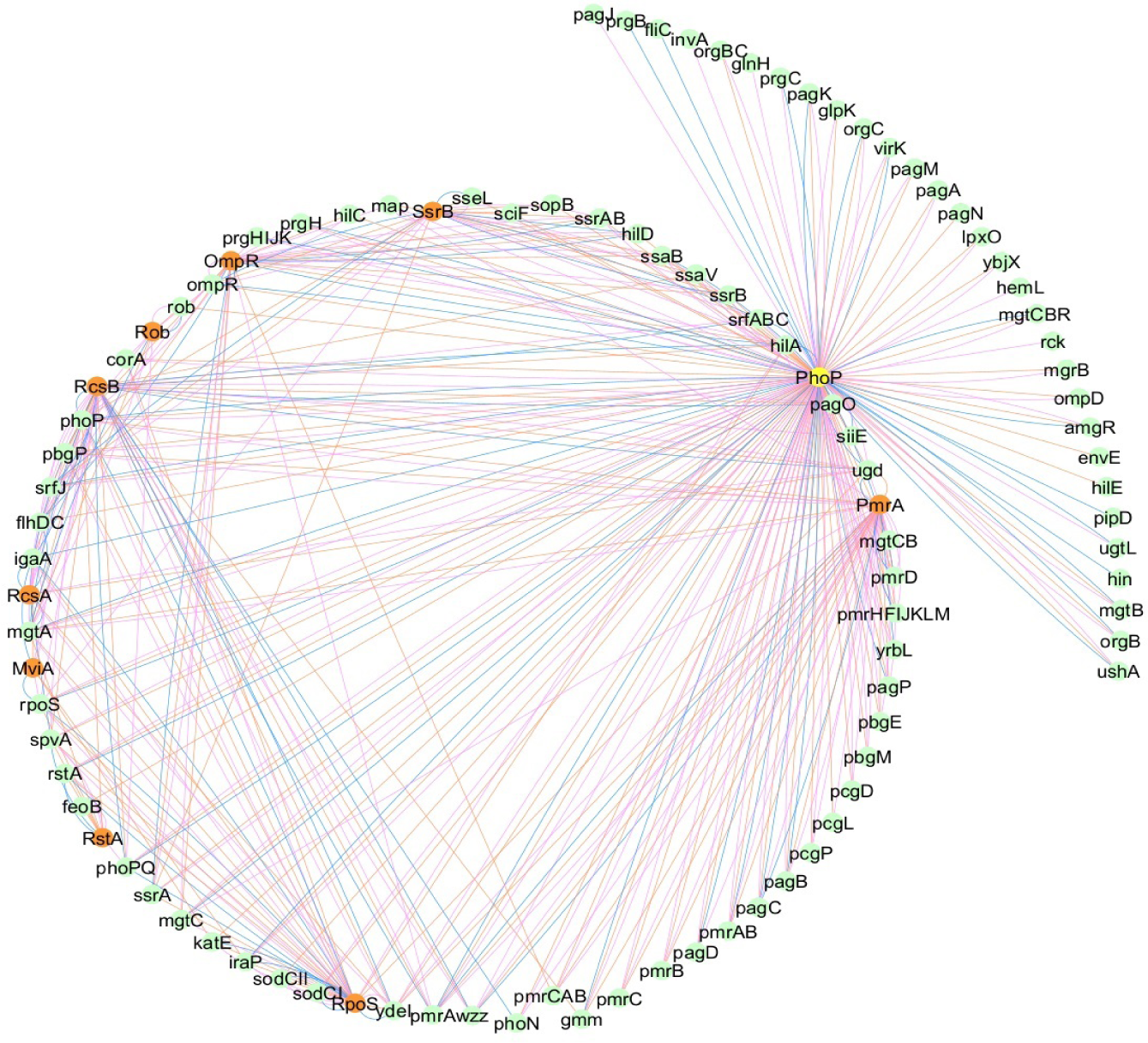
Network visualization of the PhoP community. Nodes depict Transcription factors (orange color) and regulated elements (green color). Edges depict regulatory effects: activator (pink color), repressor (blue color), and regulator (orange color).

The *no relation* category was the one that caused the most confusion to the model: seven sentences were misclassified as *activator*, five as *regulator*, and seven as *repressor* (second row in the confusion matrix). In turn, predicting the *no relation* category caused the greatest number of misclassified sentences (16 errors) despite it being the second category with more examples (second column in confusion matrix).

An examination of correct and incorrect classified sentences in the evaluation dataset was performed to extend the understanding of the predictive capabilities of our model. We confirmed that our best model learned different ways to express the regulatory effect. As expected, our model systematically predicts the correct *activator* category for sentences with the verb *activate* or some morphological variations, such as *activated, activation*. The same was observed for the *repress* and *regulate* verbs. Moreover, the model learned different lexical items to express the regulatory effect, for example, *enhance*, *stimulate*, *increase*, and *induction* for activation:

- *”The ArgP protein* ***enhances*** *the expression of the argK gene ArgP argK”* [55].
- *”Lrp* ***stimulates*** *transcription of lysP by direct binding to its control region”* [24].
- *”AraC dependent transcription initiation at the araBAD promoter is* ***increased*** *by CRP”* [56].
- *”MelR is essential for* ***induction*** *of the melAB operon that is responsible for melibiose metabolism .”* [57].

The same pattern was observed for the *repressor* category, as the model learned to associate lexical items/phrases to this category, for example, *inhibit* or *negative regulated*. For the *regulator* category we found that sentences with the verb *coregulate* was correctly classified:

- *”Thus , it appears that NarL and NarP adopt overlapping mechanisms to* ***inhibit*** *ydhY – T expression .”* [58].
- *”The treB treC operon is* ***negatively regulated*** *by TreR , whose gene treR is located upstream of treB but is not part of the operon .”* [59].
- *”Expression of acrZ is* ***coregulated*** *with acrAB and tolC by the MarA , Rob , and SoxS transcription factors .”*[60].

Regarding the misclassified sentences, we observed that our model correctly predicted some curation errors. For instance, the first following sentence (mentions of entities in boldface) was curated as *activator*, but the model predicted the true category *no relation*, as the mentions of the transcription factor GntR and the gen *gntV* does not express the regulatory interaction:

- *”This and the finding that expression of* ***gntV*** *lacZ fusion was relatively high even in the absence of cAMP ( table 4 ) may suggest that binding of* ***GntR*** *to all 3 sites slightly activates gntV expression .”* [61].

In the following sentence, the curated category was *activator* as the expression of *araC* is activated (stimulated) by CAP; however, the model predicted the category *repressor*.

- *”The expression of* ***araC*** *is repressed by its own product and stimulated by the* ***CAP*** *system ( 5 ) .”* [62].

We hypothesize that the phrase *is repressed by* was more weighted for the model prediction. This hypothesis will be in-depth explored in future work using techniques for transformer interpretability, such as those proposed by [63]. Furthermore, one would expect no error since, syntactically, the expression of *araC is… stimulated by the CAP system*” is present in the sentence, provided the model some syntactic disambiguation capabilities, which seems not to be the case. In addition, this sentence shows an interesting case, as this unveiled another way to express regulatory interactions: *auto-regulation*, which is expressed with phrases such as *is repressed by its own product*. This type of regulatory interactions can not be extracted using our strategy of passing to the BERT model the mentions of entities from a predefined list. An option to address this is to mark *its own product* being the regulatory entity.

In the following sentence, the model predicted the category *activator* for the interaction between NarL and *dmsA*, but the true effect was *repressor*.

- *”Similarly , at the FNR activated* ***NarL*** *repressed* ***dmsA*** *promoter NarL protects a large region that includes the sites for both FNR and RNA polymerase binding ( Bearson et al . , 2002 ) .”* [58].

We consider that it was due to the presence of the word *activated*, which may be more weighted for the model than the word *repressed* (again, a future study of interpretability is required). Nevertheless, the expression of the interaction is complex, as the sentence indeed has two nested regulatory interactions written with the transcription factor and the regulatory effect expressed as a past participle: *FNR activated NarL repressed dmsA*. This sentence expresses, in a compressed form, that the *dmsA* promoter is repressed by NarL and also it is activated by FNR.

A long-term challenge for biomedical relation extraction has been the co-reference. For instance, in the following sentence the model predicted the category *no relation* between SoxS and *zwf*, when the curated category was *regulator*.

- *”For instance , Rob has been shown to bind and activate the* ***zwf*** *promoter in vitro but whole cell zwf regulation cannot be activated by Rob , although the gene responds to* ***SoxS*** *and MarA ( Ariza et al . , 1995 ; Jair et al . , 1995 1996a ; b ) .”* [64].

The *regulator* effect is expressed by the phrase *the gene responds to SoxS*, where the noun phrase *the gene* refers to *zwf* (co-reference). In future work, we will review co-reference cases in our evaluation dataset to know if our BERT model learned to deal with this linguistic phenomenon. More examples of classification errors may be seen in Supplementary Table S6.

The performance scores and the analysis by categories, allow us to conclude that our model shows optimal performance on the extraction of transcriptional regulatory interaction using the sentence classification task of the regulatory effects. On the other hand, the analysis of classified examples revealed some aspects about the patterns learned by the model. The prominent pattern was that our best model weighted higher some lexical items than others, and this weighting determined the predicted category; but a deepest analysis using interpretability strategies is required.

### 4.2 Our model reconstructs a *Salmonella* TRN with high recall

To show an application of our work, we used our best LUKE model to reconstruct a TRN of *Salmonella* from 264 complete articles. The regulatory interactions of the TRN were compared with the regulatory interactions manually extracted by a curator. We found that our model was able to correctly recover and determine the type of regulation for 82% of the network (recall: 0.8217) (Table 8). Evaluating the extraction of only the transcription factor and the regulated element, the recovery of the network increases to 87% (recall: 0.8720). Nevertheless, our model extracted the double of curated interactions (1826 extracted, 909 curated), which results in a high number of apparently false positives (1079). Notice that in this case, false positives are not strictly misclassified interactions, but false positives are extracted interactions that were not in curated interactions. Evidently, these false positives include incorrect predictions, but they may include true interactions that were not present in the curated interactions. To review this aspect, we manually checked 60 randomly selected false positives interactions. Notice that one interaction may be extracted from several sentences, therefore to evaluate the interaction and its category, we reviewed the 158 sentences of the selected false positives. We found that 60% of cases were true misclassified sentences and the remaining 40% of cases were correctly predicted, corresponding to 25 new regulatory interactions (Supplementary Table S5). This finding demonstrates that our model may recover more true regulatory interactions. Some examples of true regulatory interactions that were not extracted by curation are shown in Table 9, entity mentions are in boldface.

**Table 9.** Examples of true regulatory interactions of *Salmonella* that were not extracted by curation.

| Category | TF | Regulated element | Sentence |
| --- | --- | --- | --- |
| activator | HU | *hilA* | "Thus , in contrast to *H - NS* , **HU** plays a positive role in ***hilA*** expression ." [65] |
| regulation | PhoB | *hilD* | "The regulation of ***hilD*** expression by *SirA* and **PhoB** may help mediate the regulation of *hilA* expression by these factors ." [66] |
| repressor | DeoR | *deoQ* | "The *E. coli* **DeoR** repressor was able to repress both *deoK* and ***deoQ*** expression to about the same extent as *DeoQ* ( experiment 7 versus experiments 2 and 3 ) ." [67] |

However, we must put effort into improving the precision of the model, so to deal with the false positives in future work, we propose some strategies:

1. Joining the curated datasets of *E. coli* and *Salmonella* to increase examples for fine-tuning BERT models.
2. Using the probability calculated by the model of the predicted categories to extract the most reliable ones.

The performance of our model reconstructing the majority of the curated *Salmonella* TRN shows that we may assist the manual curation of TRNs of diverse bacteria. However, a large-scale evaluation using complete networks published in databases such as RegulonDB [10] or SalmoNet [11] is an attractive opportunity to improve the method.

The network analysis revealed the relevance of the transcriptional regulatory protein PhoP, as it had the higher degree value among all transcription factors (degree: 180) (Figure 6). This finding coincides with what has been reported in the literature, as PhoP regulates 3% of the *Salmonella’s* genes. This protein belongs to the PhoP/PhoQ two-component system. PhoP has influence in virulence together with the sensor protein PhoQ and is activated by multiple signals or environmental variations, including low levels of Mg^2^, certain antimicrobial peptides and long-chain unsaturated fatty acids. PhoQ promotes the phosphorylated state of PhoP, in consequence, the phosphorylation of PhoP may have transcriptional effects on the genes and may modify the transcription in *Salmonella* [68].

As we mentioned, we obtained the community of PhoP from the TRN to describe this protein (Figure 6). We reviewed in the literature some of the genes that appeared in the community to confirm their regulation by PhoP. Next, we report these genes, if they are involved in virulence and antimicrobial resistance, and in some cases the growth conditions related to the regulation. Among these genes, it has been described that PhoP regulates *pagB* and *pmrAB*, which are responsible for the resistance to antimicrobial peptides [15]. Also, we confirmed that PhoP is a direct transcriptional activator of the *ssrB* and *rstA* genes. Other important entities present in the PhoP’s community were the *ssrb* and *rstA* genes. The SsrB regulon is modulated by the PhoP/PhoQ system [69], whereas the RstA induced by PhoP does not promote *feoB* expression at neutral pH with low magnesium content [70]. Other genes reported as being affected by the transcriptional regulatory system of PhoP/PhoQ were *mgtA* and *mgtB* [71]. Also, the genes *mtgC* and *pagC* are positively regulated by PhoP at low concentration of Mg^2^+; the PagC protein has been reported to have important implications on the interaction with the macrophage cell in the immune system [72]. Another gene activated by PhoP, which was also present in the community, was *ugtL* [73].

Within the network, there were also genes repressed by PhoP, for example, *hilC*, *hilD* and *hilE*. Extracting knowledge from these genes could contribute not only to establish their role with other genes, but also within other bacterial populations [66, 74]. We found that the mentioned conditions related to magnesium coincide with what was described in the over-representation analysis. This analysis revealed that one of the over-represented biological processes pathways was the *Magnesium ion transmembrane transport*. These results illustrate how our approach can reconstruct and expand a known TRN, further supported by the properties of the network. We also discussed the ability of the regulated genes to respond to various environmental changes, and their connection with virulence factors and drug development [75].

## 5 Limitations

In spite of our relevant findings, there is room for improvement in our work. One limitation to extract TRNs of different bacteria is that we require a list (dictionary) of transcription factors and regulated objects. This limitation will be addressed in future by coupling a Biomedical Named Entity Recognition (NER) system to identify these entities. Nevertheless, the development of strategies for biomedical NER is an active field, so we consider that the selection, evaluation and implementations of a NER system won’t be effortless [76]. A simpler strategy will be to start by generating a first version of those dictionaries from GenBank entries or genome specific databases like SalmoNet, or BioCyc for instance.

Another limitation was that our training data included only sentences with true regulatory interactions, so it was expected that our model would learn patterns associated with true interactions missing the false ones. In fact, this may be applicable to all datasets created by curation, as curation work implies recovering what is correct. For example, the first following sentence expresses a lack of regulation between YdiV and *flhDC*, but these sorts of sentences are not recovered in curation works.

- *”These average MFIs did not differ significantly between the strains ( Fig. 3B ) , indicating that* ***YdiV*** *does not regulate* ***flhDC*** *transcription .”* [77].
- *”We have also shown that* ***FadD*** , ***FliZ*** *, and* ***EnvZ*** *do not regulate* ***hilA*** *expression by modulating hilD expression .”* [66].

Our model classified this sentence from the *Salmonella* dataset as *regulator*, which was a mistake. This issue may be more complex, as there are sentences expressing a lack of regulation under specified conditions. See the second sentence, it expresses that there is no regulation between FadD, FliZ or EnvZ with *hilA*, through the modulation of *hilD* expression, so it is possible that some of these regulatory interactions occur under other conditions; in fact, this regulatory interaction (FadD, *hilA*, *regulator* ) was in the curated data of *Salmonella*.

Detecting negation, speculation, certainty and other nuances to express interactions (called *meta-knowledge dimensions*) has been in the interest of the relation extraction field for a while [78]. In future work, we will explore recent studies, especially those based on pre-trained transformer models, to detect these dimensions to improve our extraction [79, 80].

## 6 Conclusions

In this study, a BERT model to extract transcriptional regulatory interactions of bacteria from biomedical literature was fine-tuned. To the best of our knowledge, this study is the first to extract interaction between transcription factors and genes/operons using BERT architectures. The automatic or assisted extraction of regulatory interactions is of high relevance given the rich amount of knowledge present in the literature awaiting its extraction and incorporation in databases. Our best fine-tuned model was from LUKE architecture. The evaluation of our best model against curated data showed a significant performance. Our best model reconstructed a transcriptional regulatory network of *Salmonella* with high Recall. Moreover, the model was able to identify an important transcription factor, PhoP, and its community of genes. PhoP is relevant as it may have important applications in regulation studies in different bacterial populations. In future work, we will apply transformer interpretability techniques and we will improve the metrics of the model. We consider this work as a solid starting point to address the large-scale reconstruction of transcriptional regulatory networks and to explore the capabilities of BERT models for biomedical relation extraction.

## Acknowledgements

We deeply thank Dr. Julio Collado-Vides for his kind reading and accurate comments on this work, which increased its quality. We acknowledge the sources and services provided by the *Espacio de Innovacíon UNAM-HUAWEI* of the *Alianza para promover el desarrollo de capacidades digitales en México*. We acknowledge the curation team of RegulonDB and Sara Berenice Martínez-Luna for providing the initial curated data of *E. coli* and *Salmonella*, respectively. Also, we acknowledge the computational support by Víctor Del Moral and Alfredo Herńandez.

