## Supplementary Material for "Fine-tuning BERT models to extract transcriptional regulatory interactions of bacteria from biomedical literature"

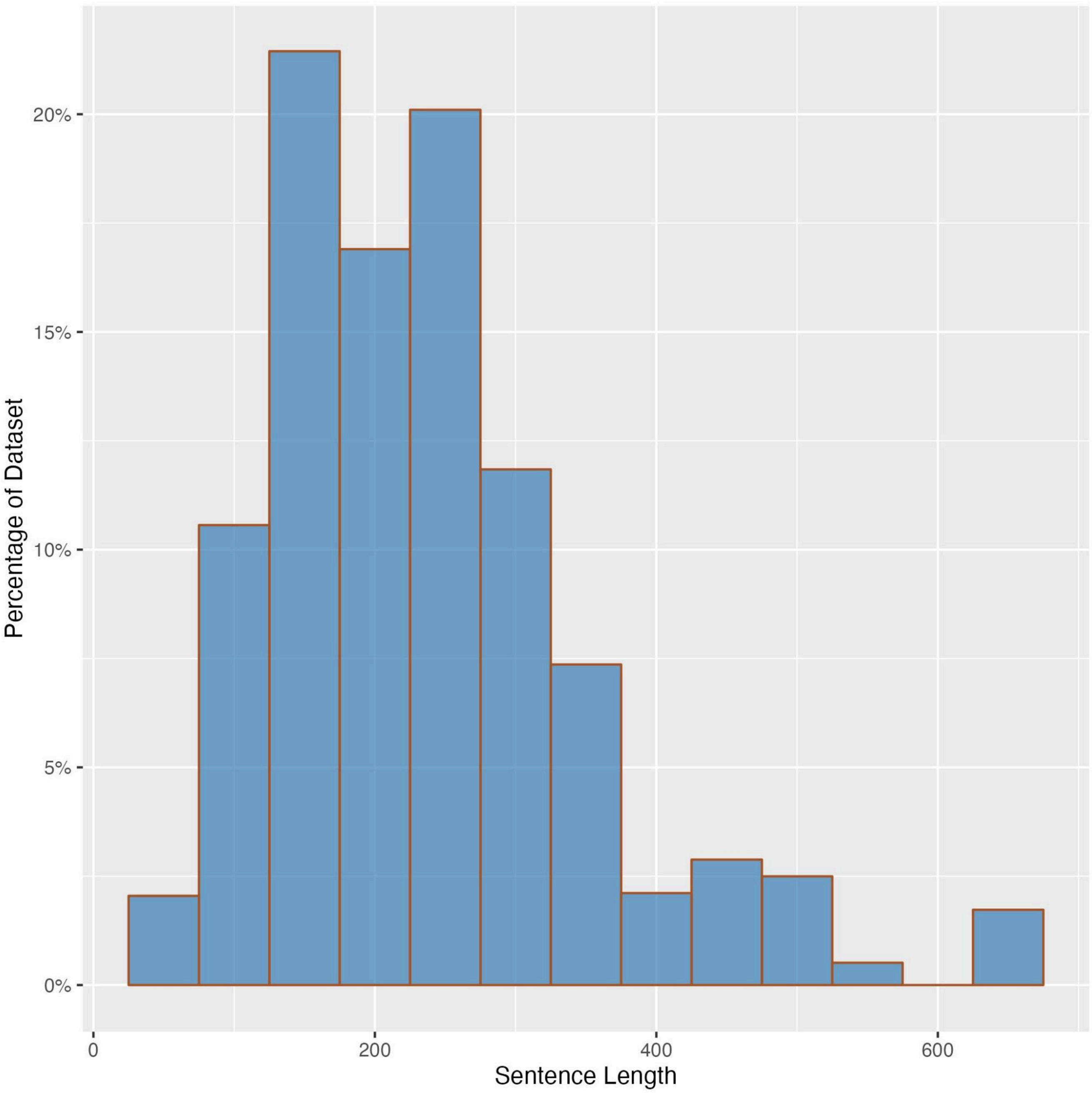


**Figure S1.** Distribution of sentence length (number of characters) in the dataset to fine-tune BERT architectures.

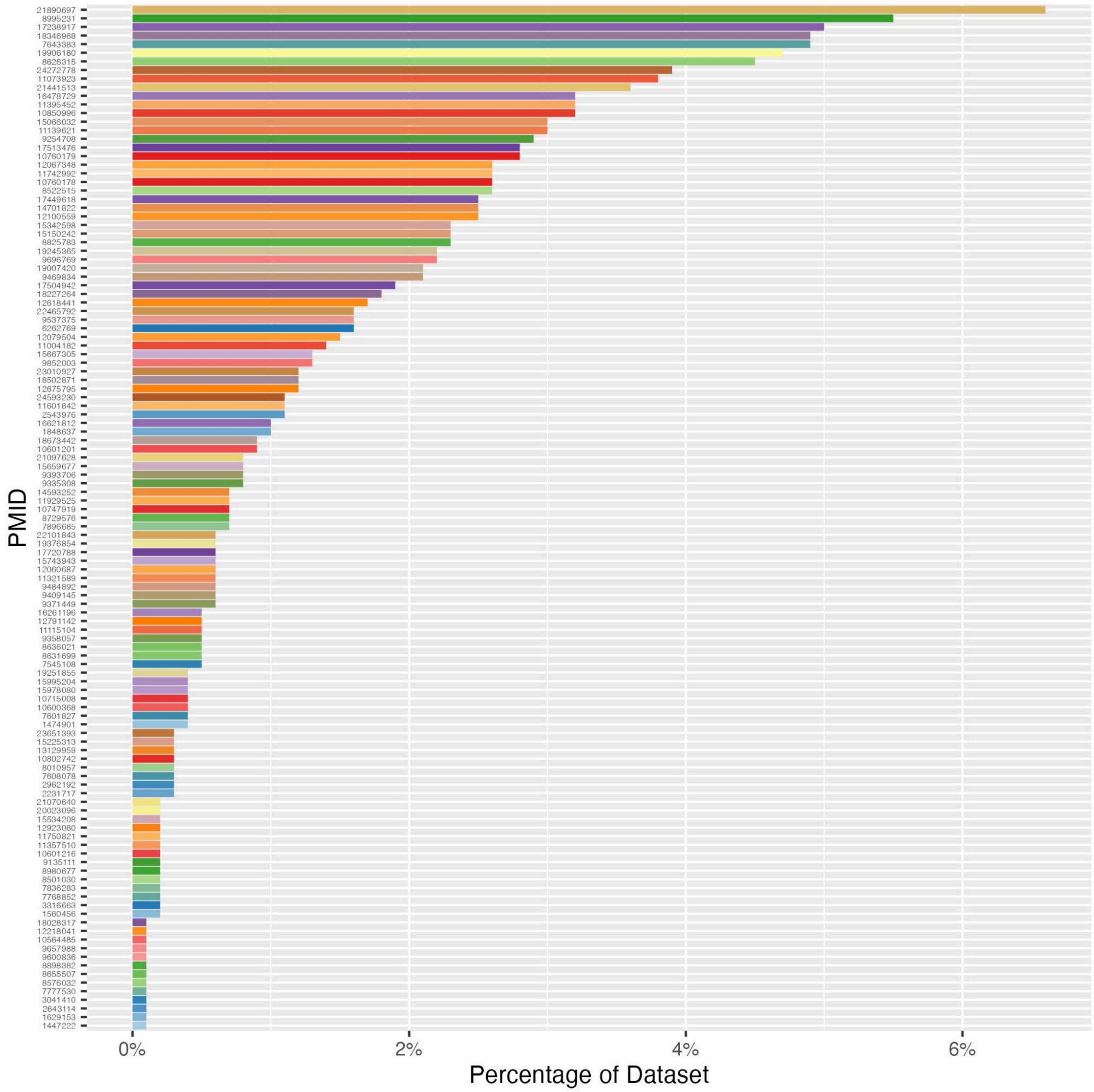


**Figure S2.** Percentage of sentences per publication (PMID) in the dataset to fine-tune BERT architectures.

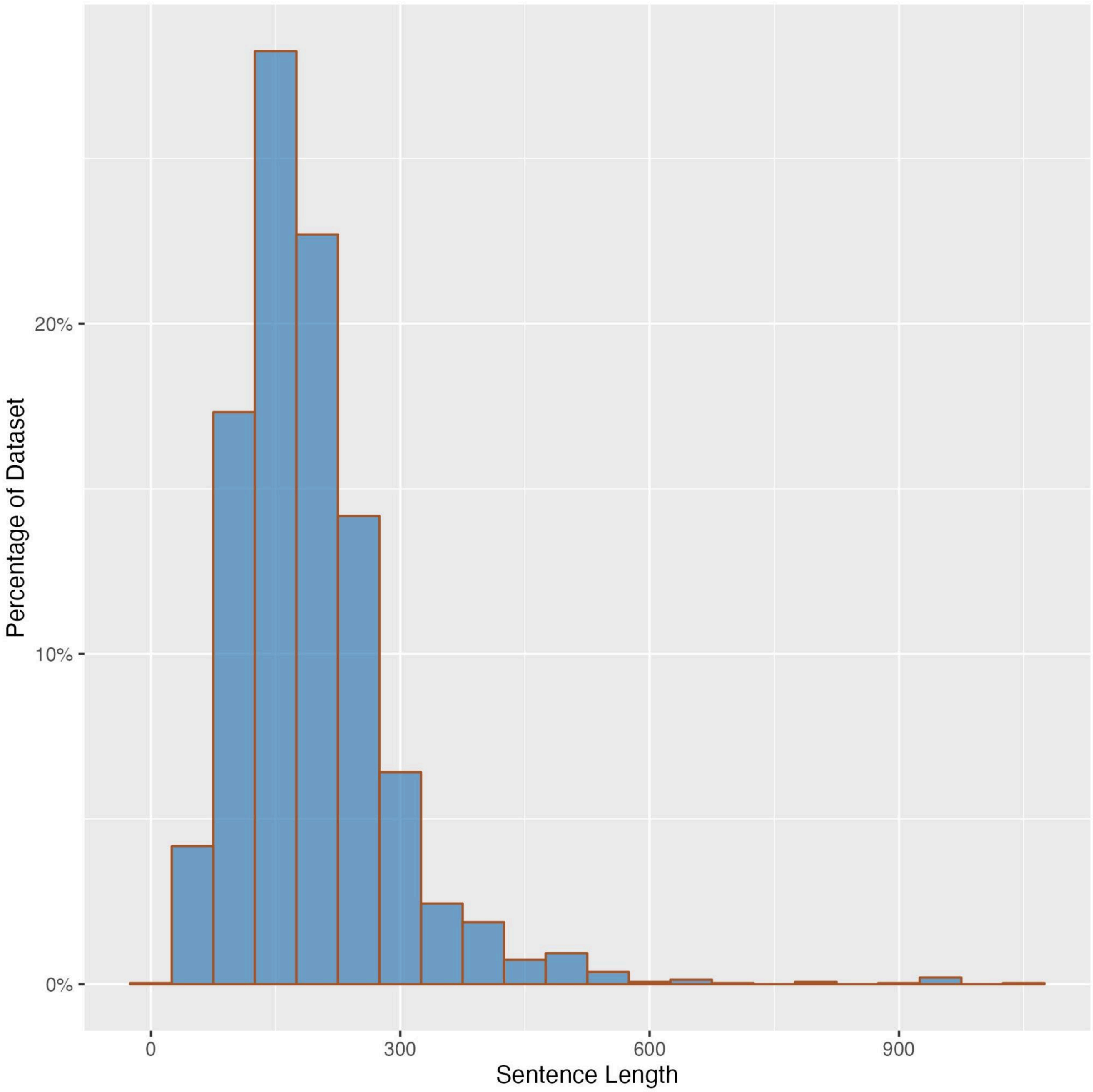


**Figure S3.** Distribution of sentence length (number of characters) in the dataset for model application.

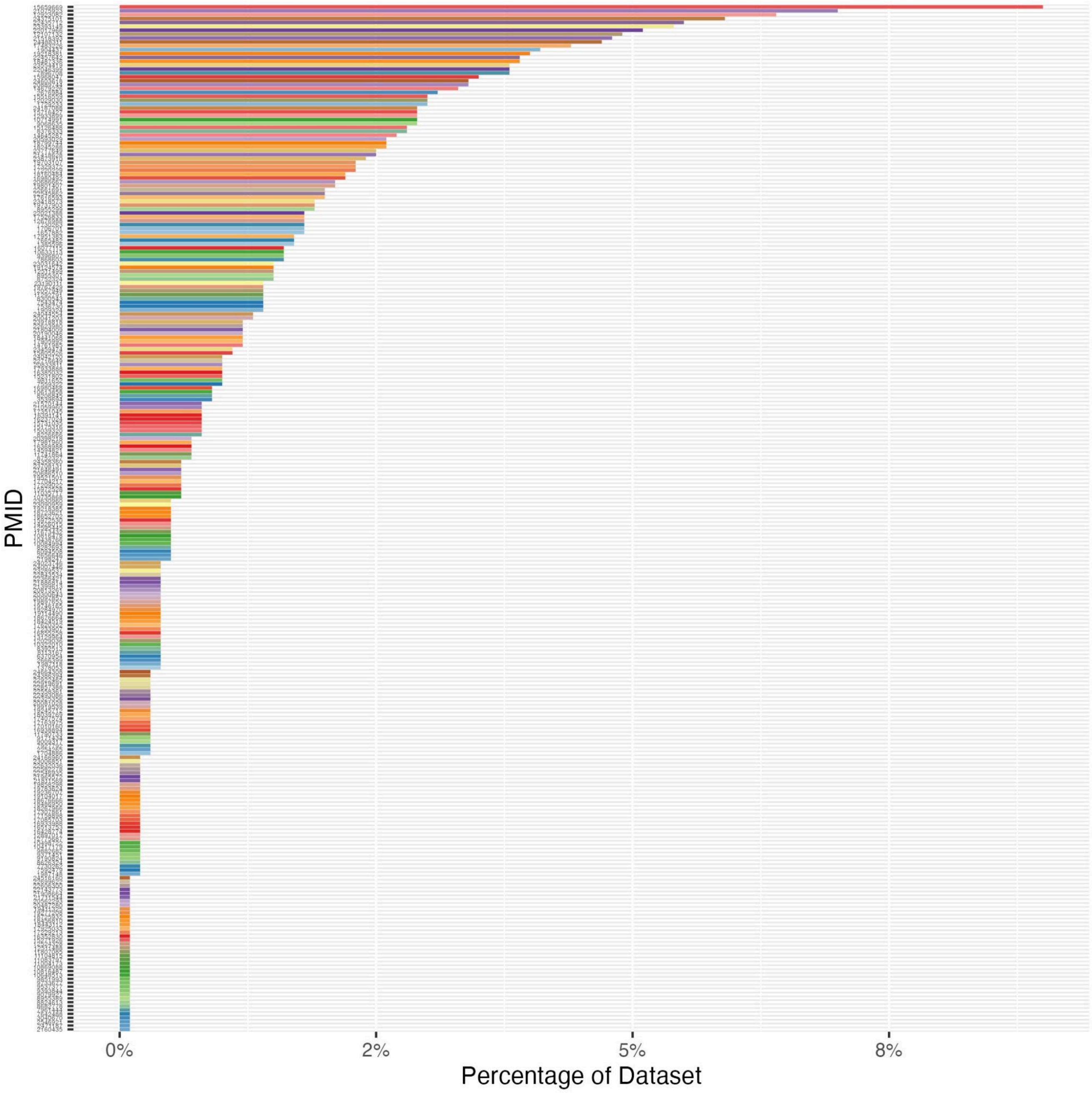


**Figure S4.** Percentage of sentences per publication (PMID) in the dataset for model application.

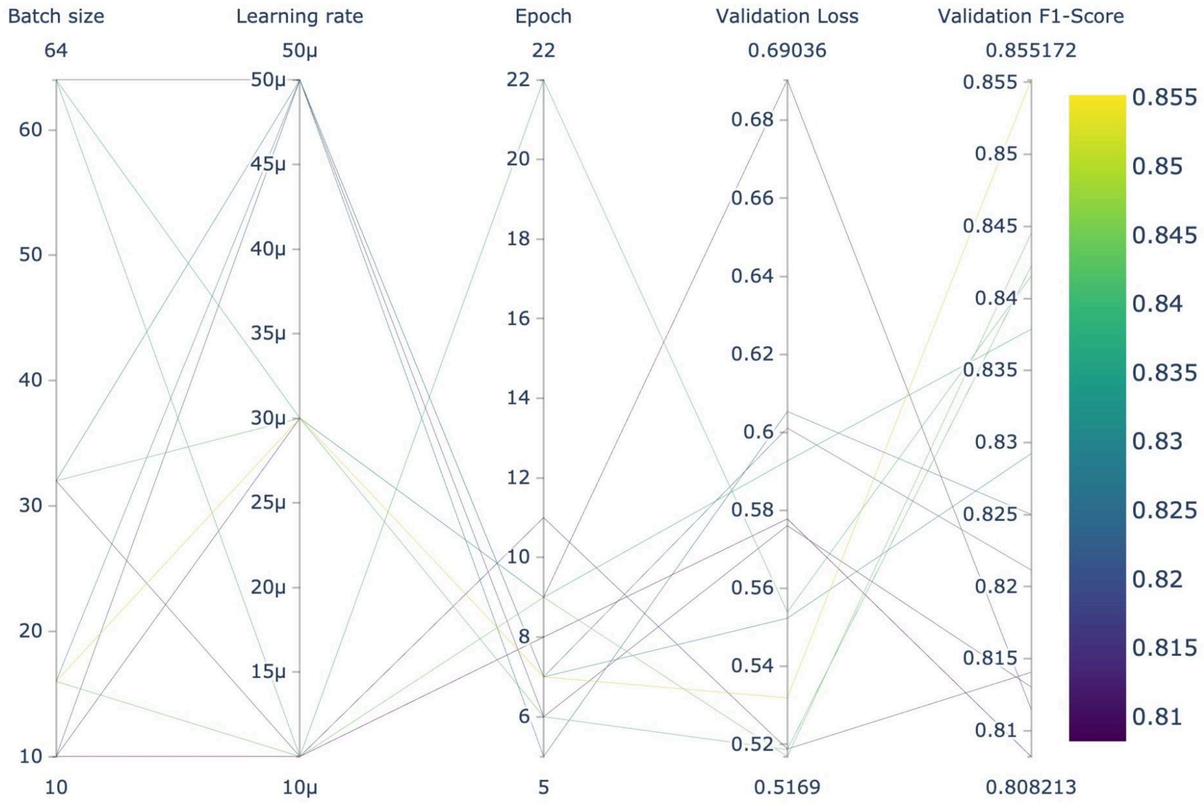


**Figure S5.** Hyper-parameter search of the twelve BERT models evaluated on the validation dataset.

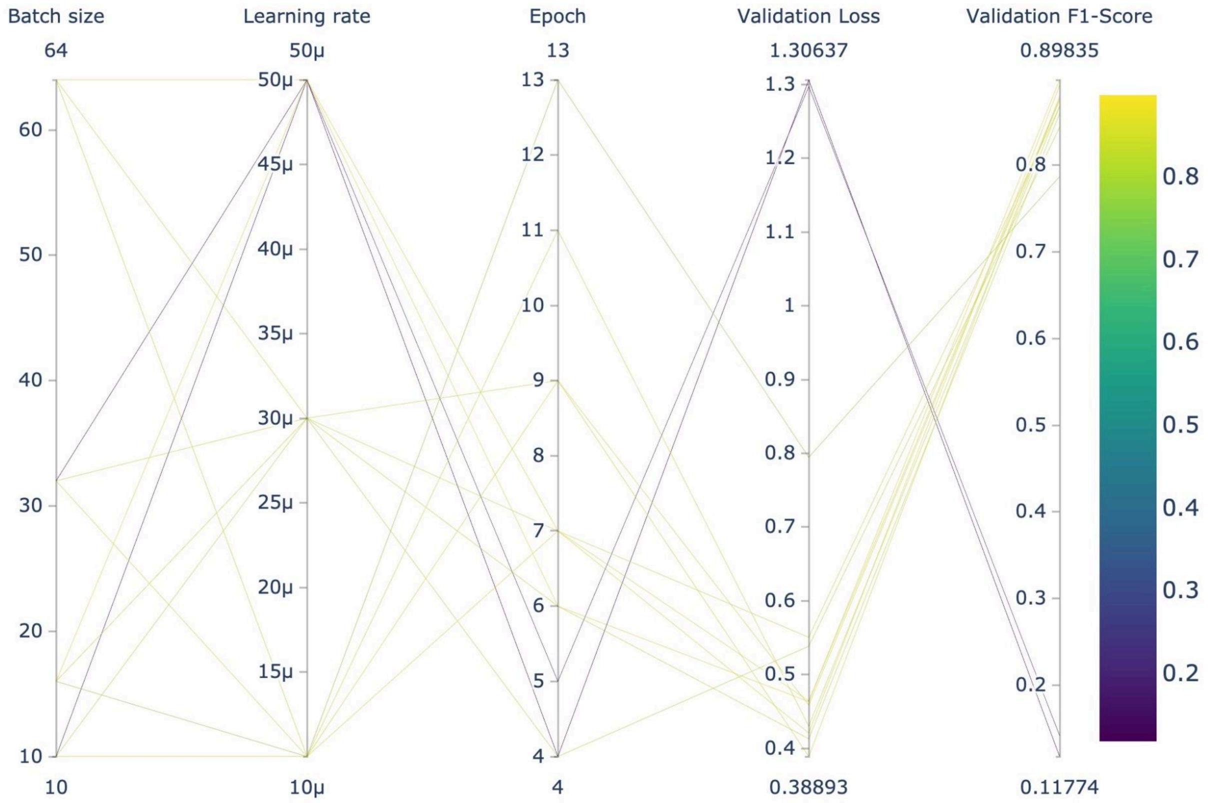


**Figure S6.** Hyper-parameter search of the twelve BioBERT models evaluated on the validation dataset.

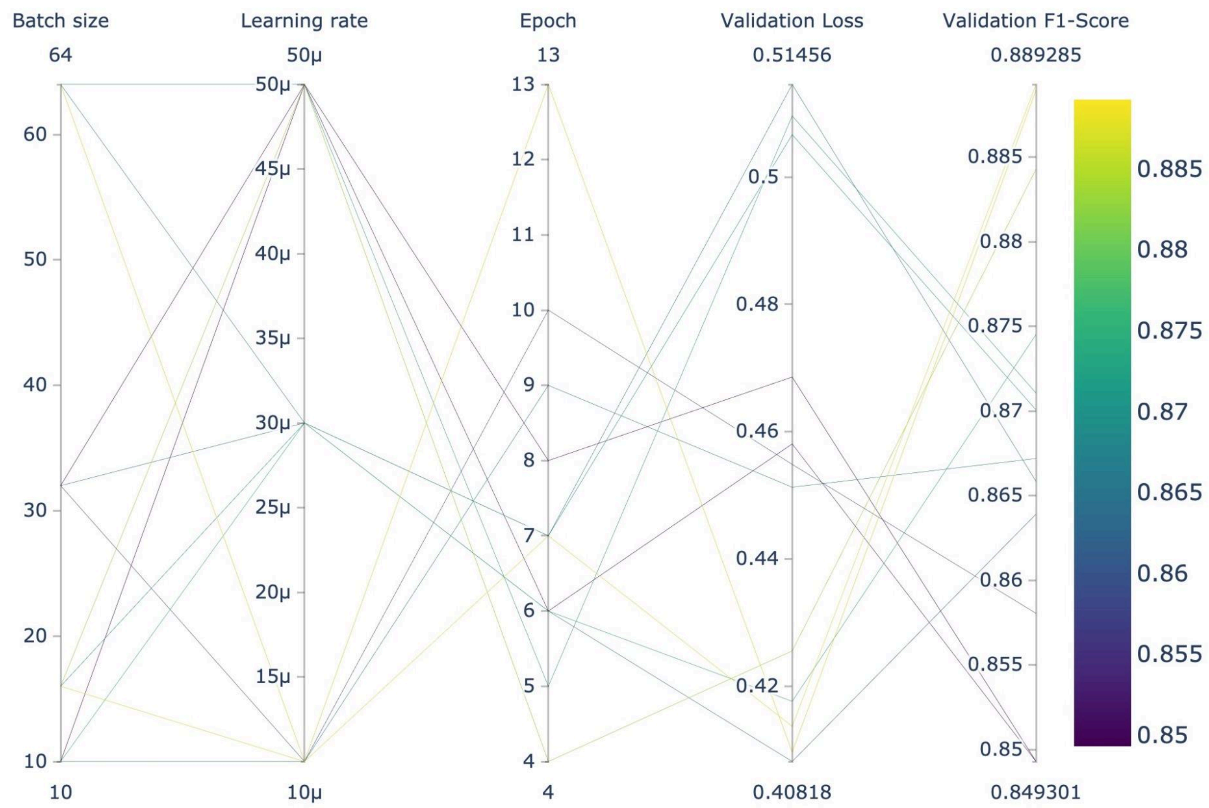


**Figure S7.** Hyper-parameter search of the twelve BioLinkBERT models evaluated on the validation dataset.

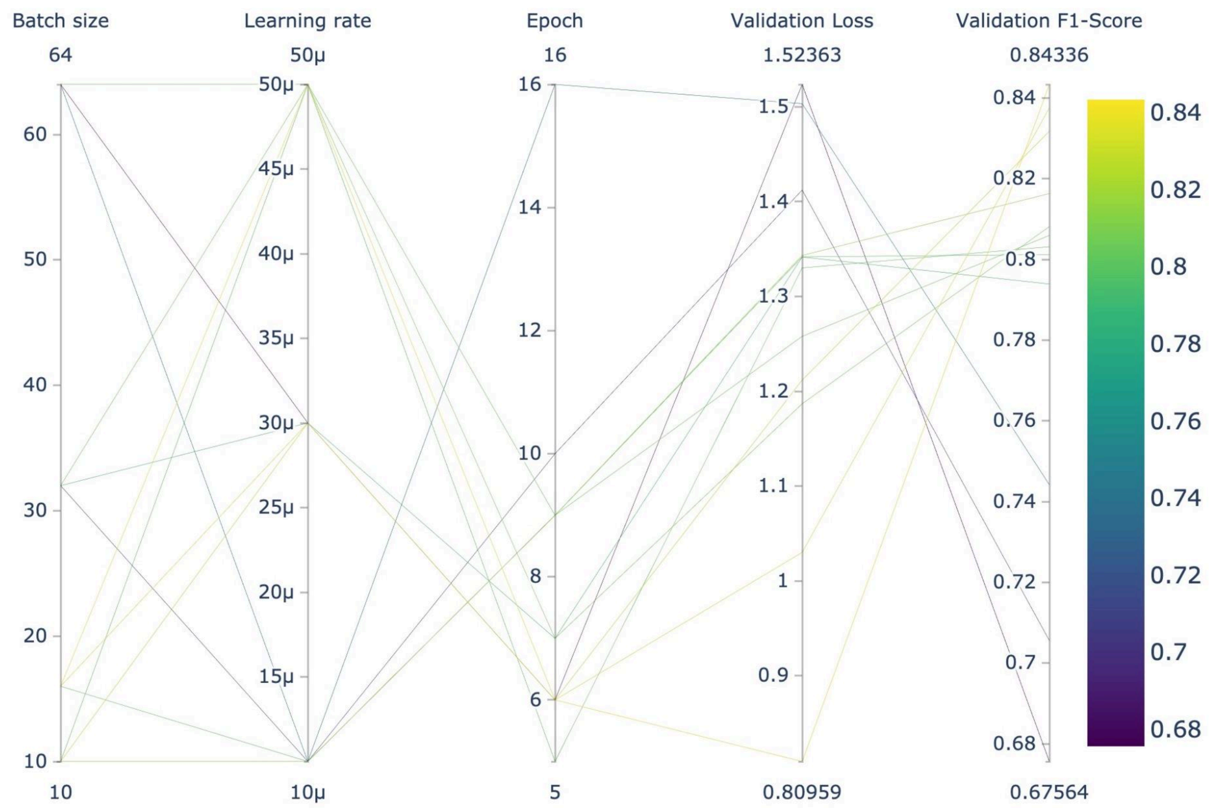


**Figure S8.** Hyper-parameter search of the twelve BioMegatron models evaluated on the validation dataset.

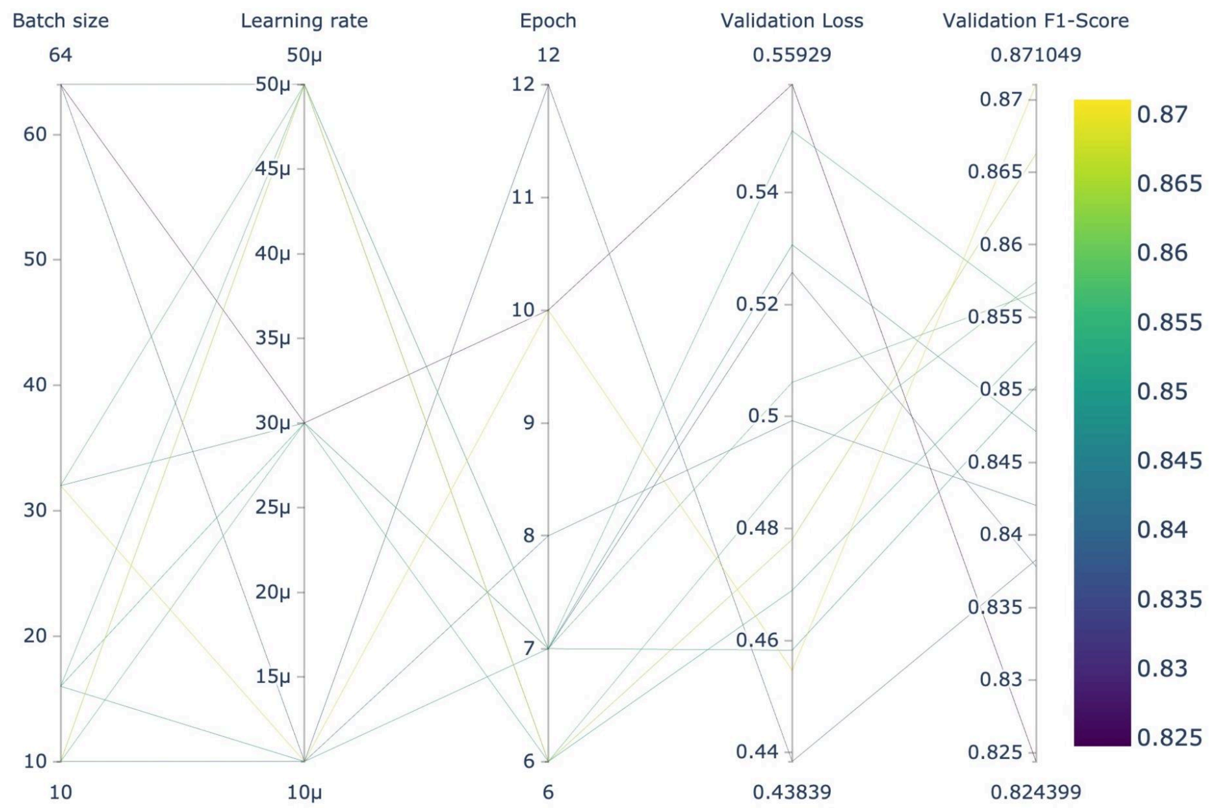


**Figure S9.** Hyper-parameter search of the twelve BioRoBERTa models evaluated on the validation dataset.

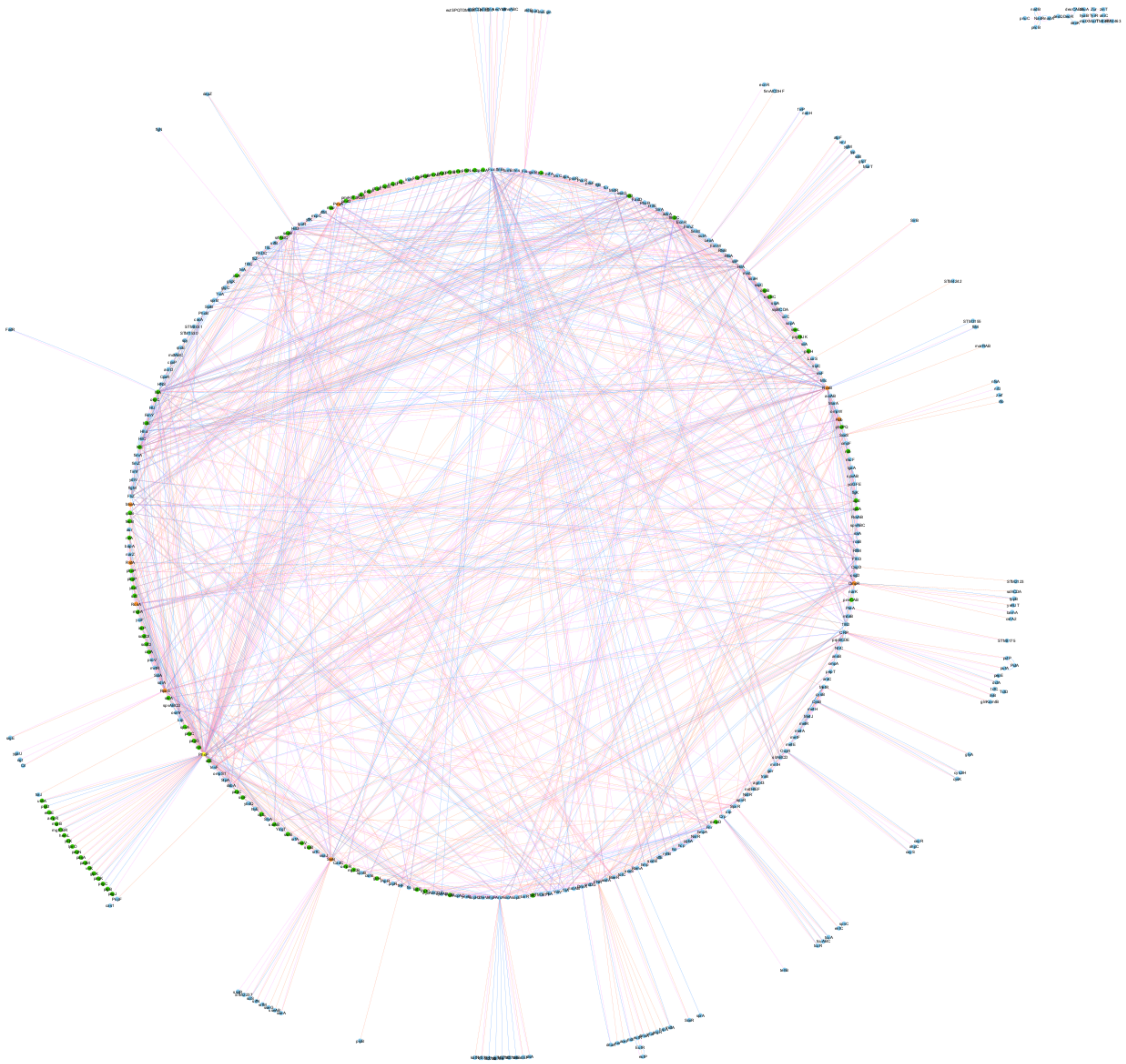

**Figure S10.** Visualization of the *Salmonella* TRN reconstructed from 264 complete articles with our best model.

**Table S1.** Comparison of characteristics between the dataset for fine-tuning and the dataset to evaluate the *Salmonella* TRN reconstruction.

| Characteristic | Dataset to find the best model | Dataset to evaluate a TRN extraction. |
|---|---|---|
| Number of sentences | 1562 | 3005 |
| % examples of Activator category | 38% | 40% |

| | | |
|---|---|---|
| % examples of Repressor category | 17% | 17% |
| % examples of Regulator category | 13% | 43% |
| Number of transcription factors | 66 | 91 |
| Number of regulated elements | 200 | 348 |
| Median of sentence length | 223 | 176 |

**Table S2.** Pre-trained BERT models downloader for our study.

| Model | Link for the implementation |
|---|---|
| BERT | https://huggingface.co/bert-base-uncased |
| BioBERT | https://huggingface.co/dmis-lab/biobert-v1.1 |
| BioLinkBERT | https://huggingface.co/michiyasunaga/BioLinkBERT-base |
| BioMegatron | https://huggingface.co/EMBO/BioMegatron345mUncased |
| BioRoBERTa | https://huggingface.co/allenai/biomed_roberta_base |
| LUKE | https://huggingface.co/studio-ousia/luke-base |

**Table S3.** Main specialized libraries and tools employed in our study.

| Library | Version | Description |
|---|---|---|
| pandas | 1.5.2 | Data analysis and manipulation |
| seaborn | 0.12.2 | Data visualization |
| matplotlib | 3.6.2 | Data visualization |
| scikit-learn | 1.0.2 | Data split and classification report |
| torch | 2.0.1 | Loading sentences to model by mini-batches and cross entropy |
| pytorch-lightning | 2.0.6 | Building deep learning models |

| | | |
|---|---|---|
| transformers | 4.29.2 | Tokenizer, transformer models with softmax classification layer and AdamW optimizer |
| wandb | 0.15.8 | Tracking training metrics in real time and sweeps for hyperparameter search |
| torchmetrics | 1.0.3 | Confusion matrix and Multiclass metrics: Precision, Recall, F1-Score, Matthew's Correlation Coefficient |

**Table S4.** Comparison of the distribution of interactions in the predicted and curated categories for Salmonella TRN reconstruction.

| Category | Number of curated interactions | Number of predicted interactions |
|---|---|---|
| Regulator | 378 (42%) | 710 (39%) |
| Activator | 321 (35%) | 661 (36%) |
| Repressor | 210 (23%) | 455 (25%) |
| Total | 909 (100%) | 1826 (100%) |

**Table S5.** The 25 regulatory interactions extracted by our model that were not in curated data.

| Transcription factor | Regulated element | Predicted category |
|---|---|---|
| HU | hilA | activator |
| PhoB | hilD | regulator |
| ArcA | rpoS | activator |
| DeoR | deoQ | repressor |
| PhoP | orgBC | activator |
| PmrA | phoP | activator |
| MviA | igaA | activator |
| RpoS | hilA | regulator |
| RcsB | srfJ | activator |
| Mlc | hilD | regulator |
| MviA | katE | activator |
| MviA | iraP | activator |
| PhoB | hilD | repressor |
| PreA | pmrCAB | activator |
| SprB | hilA | repressor |
| Fis | invE | activator |

| | | |
|---|---|---|
| FlhDC | hilD | repressor |
| InvF | slrP | activator |
| RtsA | prgHIJK | repressor |
| OxyR | metH | regulator |
| RpoS | bapA | regulator |
| CRP | putA | activator |
| PhoP | ompD | regulator |
| FimW | fimW | regulator |
| RpoS | cacA | regulator |

**Table S6.** A sample of classification errors for the evaluation dataset. Pair of mentions of entities are in boldface.

| # | Sentence | Correct category | Predicted category |
|---|---|---|---|
| 1 | *This and the finding that expression of* ***gntV*** *- lacZ fusion was relatively high even in the absence of cAMP ( table 4 ) may suggest that binding of* ***GntR*** *to all 3 sites slightly activates gntV expression .* (PMID 14593252) | activator | no_relation |
| 2 | *Both the aerobic and anaerobic expression levels of dcuB were only ca . twofold lower in the arcA mutant ( JRG3841 ) , indicating that* ***ArcA*** *plays no more than a minor role in regulating* ***dcuB*** *expression in response to oxygen ( Fig . 5B ) and that ArcA is not responsible for the FNR - independent mechanism of anaerobic activation of dcuB transcription .* (PMID 9852003) | activator | no_relation |
| 3 | *This shows that we had failed to identify MelR mutants with improved specificity for the changed KK433* ***MelR*** *- binding sequences ; mutants with specificity for the KK433 sequence would have given lower levels of activation with the wild - type* ***melAB*** *promoter .* (PMID 8010957) | activator | no_relation |
| 4 | *The expression of* ***araC*** *is repressed by its own product and stimulated by the* ***CAP*** *system ( 5 ) .* (PMID 6262769) | activator | repressor |
| 5 | ***marA*** *expression is repressed by MarR and is derepressed by the interaction of* ***MarR*** *with various phenolic compounds such as salicylate .* (PMID 12067348) | no_relation | repressor |
| 6 | *However the much stronger repression of this fusion by overproduced Mlc , compared with overproduced* ***NagC*** *, shows that the isolated* ***nagE*** *operator site has a higher affinity for Mlc than NagC .* (PMID 11139621) | no_relation | repressor |
| 7 | *Thus , the* ***melR*** *promoter is not efficiently repressed by* ***MelR*** *, and MelR is over - expressed . (PMID 18346968)* | no_relation | repressor |
| 8 | *It is unlikely that CRP binding to site 4 contributes directly to an increase in* ***rhaSR*** *expression, since transcription activation by* ***CRP*** *requires that its binding site be on the same face of the DNA as the promoter ( 6 ).* (PMID 11073923) | no_relation | activator |
| 9 | *The location of the* ***AraC*** *binding site upstream of* ***ytfQ*** *is too far upstream of the transcription start site to repress transcription by directly occluding RNAP .* (PMID 24272778) | repressor | regulator |

| | | | |
|---|---|---|---|
| 10 | *Similarly , at the FNR - activated **NarL** - repressed **dmsA** promoter NarL protects a large region that includes the sites for both FNR and RNA polymerase binding ( Bearson et al . , 2002 ) . (PMID 18227264)* | repressor | activator |
| 11 | ***MelR** carrying each of the single substitutions is less able to repress the **melR** promoter , while MelR carrying some combinations of substitutions is completely unable to repress the melR promoter .* (PMID 16621812) | repressor | no_relation |
| 12 | *On the other hand , activation of the P **araB** promoter was delayed when glucose was present , consistent with the regulation of this promoter also by **CRP** ( 6 ) .* (PMID 20023096) | regulator | activator |
| 13 | *For instance , Rob has been shown to bind and activate the **zwf** promoter in vitro but whole cell zwf regulation cannot be activated by Rob , although the gene responds to **SoxS** and MarA ( Ariza et al . , 1995 ; Jair et al . , 1995 1996a ; b ).* (PMID 12100559) | regulator | no_relation |

**Table S7.** Complete output of the Cytoscape network analysis for degree and betweenness centrality. Table sorted by degree value. The **Name** column comprises transcription factors and regulated elements.

| BetweennessCentrality | Degree | Name |
|---:|---:|---|
| 0.264210532 | 180 | PhoP |
| 0.0876723907 | 84 | HilA |
| 0.07486361657 | 82 | RcsB |
| 0.1614817061 | 77 | hilA |
| 0.113530797 | 76 | Fur |
| 0.1012641302 | 74 | RpoS |
| 0.07204096568 | 65 | OmpR |
| 0.1097447808 | 59 | flhDC |
| 0.03916432201 | 59 | HilD |
| 0.02184884007 | 54 | PmrA |
| 0.08299451706 | 53 | ArcA |
| 0.06554225205 | 53 | SsrB |
| 0.04759110046 | 48 | Fis |
| 0.03132046776 | 48 | hilD |
| 0.06611563208 | 43 | CRP |
| 0.09183332184 | 42 | FNR |
| 0.02061634812 | 37 | hilC |
| 0.01843096811 | 36 | FliZ |
| 0.01739338085 | 36 | SirA |
| 0.01964815471 | 28 | invF |

| | | |
|---|---|---|
| 0.005827495597 | 26 | HilC |
| 0.04050440911 | 25 | rpoS |
| 0.009951459224 | 25 | Fnr |
| 0.009600876761 | 25 | RtsA |
| 0.007655941718 | 25 | fliZ |
| 0.007234550616 | 24 | InvF |
| 0.04150177915 | 23 | MetR |
| 0.03797185145 | 23 | IHF |
| 0.02259035717 | 23 | OxyR |
| 0.0117093442 | 23 | flhD |
| 0.006922439377 | 23 | FlhD |
| 0.02606701296 | 22 | SoxS |
| 0.01623133146 | 22 | FlhDC |
| 0.01138165973 | 22 | hilE |
| 0.00561458934 | 22 | Hha |
| 0.005363253366 | 22 | RstA |
| 0.02524773796 | 21 | dsbA |
| 0.02006316795 | 21 | hns |
| 0.01373370077 | 21 | mgtA |
| 0.01256339968 | 20 | fliA |
| 0.02229381779 | 19 | Crp |
| 0.01099385417 | 19 | phoP |
| 0.003453977183 | 19 | RcsA |
| 0.01835957663 | 18 | ssrA |
| 0.01671387786 | 18 | ompR |
| 0.01418898433 | 18 | SpvR |
| 0.01323360235 | 18 | FimZ |
| 0.01243376244 | 18 | BaeR |
| 0.004242202975 | 18 | fimA |
| 0.01902195754 | 17 | Lrp |
| 0.01255484003 | 17 | fis |
| 0.008202901362 | 17 | siiE |
| 0.005110905787 | 17 | FliA |
| 0.01219963196 | 16 | HU |
| 0.01050608864 | 16 | SlyA |
| 0.006669483801 | 16 | Rob |

| | | |
|---|---|---|
| 0.006302546612 | 16 | NsrR |
| 0.005802490591 | 16 | TviA |
| 0.003996094602 | 16 | igaA |
| 0.003514786913 | 16 | IgaA |
| 0.009619746838 | 15 | sopB |
| 0.00880346394 | 15 | fliC |
| 0.007830922633 | 15 | ssrAB |
| 0.006393681931 | 15 | rcsB |
| 0.003377247496 | 15 | ugd |
| 0.009055272556 | 14 | spvA |
| 0.008981377462 | 14 | MarA |
| 0.003664622781 | 14 | HilE |
| 0.03226617146 | 13 | map |
| 0.01907250185 | 13 | CpxR |
| 0.009519850512 | 13 | ompW |
| 0.005050905426 | 13 | prgH |
| 0.01452648144 | 12 | sodA |
| 0.01117674642 | 12 | CysB |
| 0.002591149742 | 12 | ydeI |
| 0.001507033305 | 12 | slrP |
| 0.001264004353 | 12 | YdiV |
| 0.001143547837 | 12 | MviA |
| 0.01274358044 | 11 | phoPQ |
| 0.006036463657 | 11 | csgD |
| 0.005999962134 | 11 | hmp |
| 4.90E-04 | 11 | siiA |
| 3.47E-04 | 11 | Mlc |
| 1.85E-04 | 11 | PhoB |
| 1 | 10 | NadR |
| 0.02674667356 | 10 | spvR |
| 0.01095656506 | 10 | RamA |
| 0.009127941885 | 10 | tolC |
| 0.007716430374 | 10 | pocR |
| 0.007322740014 | 10 | crp |
| 0.00483760568 | 10 | hmpA |
| 0.002512401191 | 10 | RtsB |

| | | |
|---|---|---|
| 0.002206040403 | 10 | ssaG |
| 0.002028591565 | 10 | flhC |
| 0.001948100616 | 10 | pmrA |
| 0.001782812227 | 10 | srfJ |
| 0.001684697691 | 10 | LsrR |
| 0.001320680068 | 10 | fimZ |
| 0.00126382293 | 10 | SprB |
| 9.84E-04 | 10 | RflM |
| 7.43E-04 | 10 | rflM |
| 6.90E-04 | 10 | FimY |
| 3.49E-04 | 10 | FimW |
| 0.007596353571 | 9 | srfABC |
| 0.00750055437 | 9 | ompD |
| 0.004756013566 | 9 | prpBCDE |
| 0.004587851453 | 9 | nrdHIEF |
| 0.002591149742 | 9 | pbgP |
| 0.002470597819 | 9 | ssrB |
| 0.002346394007 | 9 | mntH |
| 8.08E-04 | 9 | sprB |
| 3.30E-04 | 9 | MntR |
| 0.009573788455 | 8 | csrB |
| 0.008924466082 | 8 | katE |
| 0.008688549093 | 8 | sulA |
| 0.008467157982 | 8 | LuxS |
| 0.00773978345 | 8 | nrdDG |
| 0.007576083336 | 8 | mgtC |
| 0.007040562077 | 8 | fur |
| 0.007027150273 | 8 | pmrCAB |
| 0.00523967046 | 8 | PreA |
| 0.00418284914 | 8 | sipC |
| 0.004069138422 | 8 | feoB |
| 0.003773010654 | 8 | rob |
| 0.001068797598 | 8 | rstA |
| 8.13E-04 | 8 | tviA |
| 7.91E-04 | 8 | fimY |
| 2.96E-04 | 8 | rcsA |

| | | |
|---|---|---|
| 6.89E-05 | 8 | ydiV |
| 0.01020113322 | 7 | LexA |
| 0.00773978345 | 7 | nrdAB |
| 0.006632985606 | 7 | sseL |
| 0.005732194772 | 7 | CsgD |
| 0.005524915344 | 7 | soxS |
| 0.005153035508 | 7 | SdiA |
| 0.004545285061 | 7 | sodB |
| 0.002846379054 | 7 | metE |
| 0.002846379054 | 7 | metH |
| 0.002346394007 | 7 | sitABCD |
| 0.001568473251 | 7 | prpR |
| 0.001465698499 | 7 | pepT |
| 0.001270123277 | 7 | CadC |
| 0.001029543486 | 7 | iraP |
| 7.70E-04 | 7 | wzz |
| 7.33E-04 | 7 | ftnB |
| 4.36E-04 | 7 | acrAB |
| 2.71E-04 | 7 | MetJ |
| 2.32E-04 | 7 | sicA |
| 2.07E-04 | 7 | PrpR |
| 1.46E-04 | 7 | NrdR |
| 1.06E-04 | 7 | pmrD |
| 0.006837972376 | 6 | dps |
| 0.005281067567 | 6 | EcnR |
| 0.005224604743 | 6 | mgtCB |
| 0.005144915887 | 6 | sirC |
| 0.005089058524 | 6 | EutR |
| 0.002958552714 | 6 | spvB |
| 0.002660838647 | 6 | ackA |
| 0.002296870093 | 6 | mntR |
| 0.00194693085 | 6 | RcsAB |
| 0.001604936223 | 6 | ompF |
| 0.001573421396 | 6 | ssaB |
| 0.001254151858 | 6 | sodCII |
| 7.21E-04 | 6 | fljB |

| | | |
|---|---|---|
| 6.64E-04 | 6 | fimW |
| 6.15E-04 | 6 | pagC |
| 5.13E-04 | 6 | bapA |
| 5.05E-04 | 6 | StpA |
| 2.27E-04 | 6 | PocR |
| 1.10E-04 | 6 | yciGFE |
| 7.69E-05 | 6 | narZ |
| 5.72E-05 | 6 | FadD |
| 0.005369733698 | 5 | misL |
| 0.00379083757 | 5 | pmrC |
| 0.002538038116 | 5 | putP |
| 0.002413954309 | 5 | prgHIJK |
| 0.00149240067 | 5 | corA |
| 0.001442818118 | 5 | ssaV |
| 0.001254151858 | 5 | sodCI |
| 0.001004084159 | 5 | orgB |
| 9.30E-04 | 5 | cydAB |
| 7.70E-04 | 5 | gmm |
| 6.12E-04 | 5 | asr |
| 3.65E-04 | 5 | pduF |
| 3.38E-04 | 5 | ssaH |
| 1.47E-04 | 5 | yneB |
| 1.26E-04 | 5 | sopE |
| 1.25E-04 | 5 | PhoR |
| 1.06E-04 | 5 | pagB |
| 1.06E-04 | 5 | pagP |
| 1.06E-04 | 5 | pmrAB |
| 1.06E-04 | 5 | pmrHFIJKLM |
| 7.66E-05 | 5 | sopA |
| 7.59E-05 | 5 | flgM |
| 3.22E-05 | 5 | ytfK |
| 6.49E-06 | 5 | PutA |
| 1 | 4 | DeoR |
| 0.01015215246 | 4 | eutR |
| 0.00810514144 | 4 | glnH |

| | | |
|---|---|---|
| 0.00521248213 | 4 | NtrC |
| 0.005091654983 | 4 | DnaA |
| 0.003817032102 | 4 | sdiA |
| 0.002538038116 | 4 | putA |
| 0.002244390531 | 4 | ytfE |
| 0.001829826797 | 4 | mgtB |
| 0.001592635758 | 4 | sspH2 |
| 0.001511527588 | 4 | micF |
| 0.001344674594 | 4 | invA |
| 0.001305794646 | 4 | spvABCD |
| 0.00129402544 | 4 | iroA |
| 0.00120798138 | 4 | flgK |
| 0.001016230656 | 4 | agn43 |
| 9.30E-04 | 4 | cyoABCDE |
| 8.78E-04 | 4 | ansB |
| 8.19E-04 | 4 | metA |
| 8.19E-04 | 4 | metF |
| 8.19E-04 | 4 | metR |
| 7.70E-04 | 4 | phoN |
| 7.30E-04 | 4 | spvABC |
| 6.83E-04 | 4 | cysB |
| 6.53E-04 | 4 | ryhB |
| 5.09E-04 | 4 | cacA |
| 4.78E-04 | 4 | mcpC |
| 4.55E-04 | 4 | pagO |
| 3.86E-04 | 4 | orgC |
| 3.30E-04 | 4 | ompS1 |
| 2.84E-04 | 4 | ugtL |
| 1.61E-04 | 4 | YdgT |
| 1.06E-04 | 4 | pagD |
| 1.06E-04 | 4 | pbgE |
| 1.06E-04 | 4 | pmrB |
| 3.22E-05 | 4 | lpxR |
| 1 | 3 | Zur |
| 0.006360150469 | 3 | rck |
| 0.002011697528 | 3 | trxB |

| | | |
|---|---|---|
| 0.001176959954 | 3 | sciF |
| 0.001056385123 | 3 | ahpF |
| 0.001004084159 | 3 | orgBC |
| 8.97E-04 | 3 | araC |
| 7.33E-04 | 3 | fumB |
| 7.28E-04 | 3 | rpoE |
| 6.80E-04 | 3 | yciF |
| 6.53E-04 | 3 | hcp |
| 6.53E-04 | 3 | hcr |
| 6.53E-04 | 3 | ygbA |
| 6.47E-04 | 3 | srfB |
| 6.41E-04 | 3 | osmY |
| 6.10E-04 | 3 | srfA |
| 5.96E-04 | 3 | hin |
| 5.09E-04 | 3 | cpxP |
| 5.04E-04 | 3 | mdaB |
| 3.92E-04 | 3 | invC |
| 3.49E-04 | 3 | STM3611 |
| 1.37E-04 | 3 | proV |
| 1.08E-04 | 3 | STM3031 |
| 1.08E-04 | 3 | acrD |
| 1.08E-04 | 3 | mdtABC |
| 1.06E-04 | 3 | pbgM |
| 1.06E-04 | 3 | pcgD |
| 1.06E-04 | 3 | pcgP |
| 1.06E-04 | 3 | yrbL |
| 6.40E-05 | 3 | spvAB |
| 5.59E-05 | 3 | sspC |
| 0 | 3 | mdtA |
| 0 | 3 | dctA |
| 0 | 3 | tcuA |
| 0 | 3 | tcuABC |
| 0 | 3 | tcuR |
| 0 | 3 | cysJIH |
| 0 | 3 | cysK |
| 0 | 3 | eutP |

| | | |
|---|---|---|
| 0 | 3 | ftnA |
| 0 | 3 | lpp |
| 0 | 3 | ygiM |
| 0 | 3 | dapZ |
| 0 | 3 | MarT |
| 0 | 3 | nadA |
| 0 | 3 | nadB |
| 0 | 3 | pncB |
| 0 | 3 | tppB |
| 0 | 3 | oxyR |
| 0 | 3 | oxyS |
| 0 | 3 | amgR |
| 0 | 3 | mgtCBR |
| 0 | 3 | pagK |
| 0 | 3 | ushA |
| 0 | 3 | virK |
| 0 | 3 | STM4463 |
| 0 | 3 | STM4467 |
| 0 | 3 | sciS |
| 1 | 2 | TyrR |
| 0.005089058524 | 2 | tctC |
| 0.005089058524 | 2 | PspF |
| 0.002011697528 | 2 | gor |
| 7.41E-04 | 2 | ssaJ |
| 7.22E-04 | 2 | ompA |
| 5.48E-04 | 2 | cheV |
| 5.48E-04 | 2 | mcpAC |
| 5.09E-04 | 2 | spy |
| 4.46E-04 | 2 | hlyE |
| 3.32E-04 | 2 | srfC |
| 2.87E-04 | 2 | gyrB |
| 1.88E-04 | 2 | nrfA |
| 1.29E-04 | 2 | HNS |
| 1.13E-04 | 2 | fliL |
| 1.09E-04 | 2 | narK |
| 1.08E-04 | 2 | STM1530 |

| | | |
|---|---|---|
| 1.06E-04 | 2 | pcgL |
| 7.05E-05 | 2 | ybdQ |
| 5.02E-05 | 2 | glgC |
| 5.02E-05 | 2 | glgX |
| 2.80E-05 | 2 | sipA |
| 2.80E-05 | 2 | sipBCDA |
| 0 | 2 | STM3784 |
| 0 | 2 | nrdEF |
| 0 | 2 | slsA |
| 0 | 2 | ilvB |
| 0 | 2 | deoCABD |
| 0 | 2 | dnaA |
| 0 | 2 | dmsA |
| 0 | 2 | ndh |
| 0 | 2 | FadR |
| 0 | 2 | invE |
| 0 | 2 | spaO |
| 0 | 2 | secY |
| 0 | 2 | atpF |
| 0 | 2 | glpT |
| 0 | 2 | finP |
| 0 | 2 | nanH |
| 0 | 2 | MalT |
| 0 | 2 | malX |
| 0 | 2 | glyA |
| 0 | 2 | glnKamtB |
| 0 | 2 | glpK |
| 0 | 2 | lpxO |
| 0 | 2 | mgrB |
| 0 | 2 | pagA |
| 0 | 2 | pagM |
| 0 | 2 | prgC |
| 0 | 2 | ybjX |
| 0 | 2 | STM3175 |
| 0 | 2 | fliM |
| 0 | 2 | Crl |

| | | |
|---|---|---|
| 0 | 2 | srgE |
| 0 | 2 | SirB |
| 0 | 2 | SoxR |
| 0 | 2 | entC |
| 0 | 2 | srfN |
| 0 | 2 | sseA |
| 0 | 2 | zinT |
| 0 | 1 | STM2786 |
| 0 | 1 | STM2787 |
| 0 | 1 | STM2788 |
| 0 | 1 | STM2789 |
| 0 | 1 | katN |
| 0 | 1 | tctE |
| 0 | 1 | pepE |
| 0 | 1 | STM2123 |
| 0 | 1 | deoK |
| 0 | 1 | deoQ |
| 0 | 1 | ecnR |
| 0 | 1 | aceF |
| 0 | 1 | aer |
| 0 | 1 | hrpA |
| 0 | 1 | ycgR |
| 0 | 1 | ynaF |
| 0 | 1 | fimAICDHF |
| 0 | 1 | aldB |
| 0 | 1 | gin |
| 0 | 1 | flgN |
| 0 | 1 | bfr |
| 0 | 1 | dmsABC |
| 0 | 1 | eutSPQTDMEJGHABCLK |
| 0 | 1 | iroBCDE |
| 0 | 1 | ydiE |
| 0 | 1 | isrJ |
| 0 | 1 | siiB |

| | | |
|---|---|---|
| 0 | 1 | pipB |
| 0 | 1 | traJ |
| 0 | 1 | STM4242 |
| 0 | 1 | marRAB |
| 0 | 1 | pnuC |
| 0 | 1 | tehB |
| 0 | 1 | bamA |
| 0 | 1 | cstA2 |
| 0 | 1 | sdhCDA |
| 0 | 1 | yehUT |
| 0 | 1 | ahpC |
| 0 | 1 | envE |
| 0 | 1 | hemL |
| 0 | 1 | pagJ |
| 0 | 1 | pagN |
| 0 | 1 | pipD |
| 0 | 1 | prgB |
| 0 | 1 | cas1 |
| 0 | 1 | sprA |
| 0 | 1 | STM3155 |
| 0 | 1 | ogt |
| 0 | 1 | ygaU |
| 0 | 1 | nfo |
| 0 | 1 | nfsA |
| 0 | 1 | nifJ |
| 0 | 1 | zwf |
| 0 | 1 | spvC |
| 0 | 1 | STM2237 |
| 0 | 1 | srfM |
| 0 | 1 | ssaR |
| 0 | 1 | sseAB |
| 0 | 1 | sseG |
| 0 | 1 | TctD |
| 0 | 1 | aniC |
| 0 | 1 | hyaB |
| 0 | 1 | znuA |
